# Phylogenetic turnover during subtropical forest succession across environmental and phylogenetic scales

**DOI:** 10.1101/162727

**Authors:** Oliver Purschke, Stefan G. Michalski, Helge Bruelheide, Walter Durka

## Abstract

- Although spatial and temporal patterns of phylogenetic community structure during succession are inherently interlinked and assembly processes vary with environmental and phylogenetic scale, successional studies of community assembly have yet to integrate spatial and temporal components of community structure, while accounting for scaling issues. To gain insight into the processes that generate biodiversity after disturbance, we combine analyses of spatial and temporal phylogenetic turnover across phylogenetic scales, accounting for covariation with environmental differences.
- We compared phylogenetic turnover, at the species-and individual-level, within and between five successional stages, representing woody plant communities in a subtropical forest chronosequence. We decomposed turnover at different phylogenetic depths and assessed its covariation with between-plot abiotic differences.
- Phylogenetic turnover between stages was low relative to species turnover and was not explained by abiotic differences. However, within the late successional stages, there was high presence/absence-based turnover (clustering) that occurred deep in the phylogeny and covaried with environmental differentiation.
- Our results support a deterministic model of community assembly where (i) phylogenetic composition is constrained through successional time, but (ii) towards late succession, species sorting into preferred habitats according to niche traits that are conserved deep in phylogeny, becomes increasingly important.

## Introduction

A better understanding of the processes that generate biodiversity during succession after disturbance is needed for more accurate predictions of ecosystem responses to future disturbance events (Garnier *et al.,* 2004; Dornelas, 2010). Community assembly during succession may be driven by deterministic (biotic and abiotic filtering) as well as stochastic processes (Keddy, 1992; Fukami *et al.,* 2005) that are often inferred using trait-based approaches (Bazzaz, 1979; Shipley *et al.,* 2006). However, the traits involved in assembly processes are *a priori* unknown and, particularly in species rich systems, it is difficult to choose and measure the most relevant traits. In communities with broad taxonomic sampling, such as hyper-diverse tropical plant communities, closely related species often share similar functional characteristics (Swenson, 2013), resulting from phylogenetic niche conservatism (Losos, 2008). In such systems, phylogenetic relatedness between species is often used as a proxy for overall trait similarity as it potentially integrates more trait information than a limited set of measurable traits (Pavoine & Bonsall, 2011; Mouquet *et al.,* 2012). Several studies have quantified spatial or temporal patterns of phylogenetic relatedness throughout succession, either by testing for non-random patterns of relatedness within successional stages (Letcher, 2010; Ding *et al.,* 2012) or by examining whether the observed temporal phylogenetic turnover between stages differed from the expected phylogenetic turnover, given the level of species turnover (Swenson *et al.,* 2012, Letten *et al.,* 2014). However, purely temporal approaches, that focus on phylogenetic turnover *between* stages, do not allow to evaluate whether non-random patterns of temporal phylogenetic turnover are simply a reflection of spatial turnover between sites belonging to the same successional stage (see Purschke *et al.,* 2013). In contrast, approaches that focus on spatial patterns of phylogenetic relatedness *within* successional stages only allow for inferences about assembly processes that act at a particular successional stage. Because spatial and temporal patterns of community composition are inherently interlinked (Preston, 1960; White *et al.,* 2010), studies based on partial analysis of either spatial or temporal patterns of community phylogenetic structure during succession will only give limited insight into the temporal dynamics of assembly processes.

Hardy and Senterre (2007) proposed a framework that allows to test the spatial phylogenetic structure of communities, based on the extent to which species within sites are more, or less, related to each other than to species from different sites. If species that co-occur within a site are more related to each other than to species from different sites, phylogenetic turnover between sites is high, which is referred to as spatial phylogenetic clustering. Such high phylogenetic turnover is usually interpreted as a signature of abiotic filtering where distinct groups of closely related, and functionally similar, species are differentially selected in sites that differ in their environmental conditions (Baraloto *et al.,* 2012). Alternatively, phylogenetic clustering may reflect the exclusion of competitively inferior species, i.e. competitive hierarchies, if the traits conferring competitive dominance are phylogenetically conserved (Mayfield & Levine, 2010). In contrast, if species within sites are phylogenetically less related than species from different sites, phylogenetic turnover between sites is low, which is referred to as spatial phylogenetic overdispersion. This pattern is often interpreted as result of biotic filtering because of negative interactions due to limiting similarity competition between closely related species, but could also indicate abiotic filtering in case of convergent evolution of important niche traits (Cavender-Bares *et al.,* 2004). Because the Hardy & Senterre (2007) framework expresses community differentiation between sites, it can also be applied to pairs of communities at different successional stages (see Purschke *et al.,* 2013), allowing to compare spatial and temporal patterns of community differentiation within a consistent framework.

Despite the promise of combining spatial and temporal components of phylogenetic turnover to gain insight into assembly processes, there remain several difficulties with interpreting community phylogenetic structure. One main problem is that patterns of phylogenetic relatedness within communities and conclusions about assembly processes are highly scale-dependent (Swenson *et al.,* 2007; Graham *et al.*, 2016). For instance, patterns of phylogenetic overdispersion will only be detectable at small environmental, spatial and phylogenetic scales (i.e. between closely related species close to tips of the phylogeny, see Parmentier *et al.,* 2014). In contrast, phylogenetic clustering, resulting from abiotic filtering, has mainly been demonstrated over steep to moderate ecological gradients and at large phylogenetic scales, i.e. deep in the phylogeny (Cavender-Bares *et al.,* 2006). In addition, Hardy & Senterre (2007) pointed out that if such opposing assembly mechanisms, like overdispersion and clustering, act simultaneously at different phylogenetic scales, they may cancel out each other, resulting in an overall random phylogenetic structure. To address this phylogenetic scaling issue, phylogenetic structure can be assessed at different depths in the phylogenetic tree (Hardy & Senterre, 2007; Cavender-Bares & Reich, 2012). The issue of environmental scaling may be accounted for by assessing the extent to which phylogenetic turnover is explained by environmental differences between sites (e.g. Hardy *et al.,* 2012).

Finally, inferences about assembly processes may be influenced by the level of biological organization considered in the analysis, i.e. whether phylogenetic structure is assessed on the level of species or individuals, respectively, giving more weight to rare or dominant species (Helmus *et al.,* 2007; Lozupone *et al.,* 2007). The joint use of abundance- and presence/absence-based indices allows to detect the relative importance of shifts in species abundances vs. changes in composition, and hence will be critical to understand the processes underlying community assembly (Vellend *et al.,* 2011).

In the context of succession, theory predicts that in early succession, disturbance acts as an environmental filter selecting for closely related species and that biotic filtering will become more important over time, selecting for more distantly related species in late succession (Cornell & Slatyer, 1977). While a number of studies found support for this hypothesis (e.g. Letcher, 2010; Whitfield *et al.,* 2012; Purschke *et al.,* 2013), a few recent studies detected an increase in phylogenetic relatedness during succession, and suggested that hierarchical competition and/or environmental filtering become more important during succession (e.g. Uriarte *et al.,* 2010; Kunstler *et al.,* 2012; Letten *et al.,* 2014; Buzzard *et al.,* 2015). However, existing studies of phylogenetic community structure (i) were usually based on metrics of phylogenetic structure that integrate across the whole phylogeny, and therefore did not allow for the possibility that assembly processes will only be detectable at particular phylogenetic scales, (ii) did not include information on environmental differentiation between sites, or (iii) focused either on spatial or temporal components of community change. To gain more accurate insights into the processes that underlie community assembly during succession after disturbance, there is therefore a need for integrative studies that account for phylogenetic community structure at different phylogenetic scales and that compare spatial and temporal turnover components in conjunction with environmental differentiation between sites. If, for example, abiotic filtering along an environmental gradient is the predominant process shaping communities at the beginning of succession and there is phylogenetic conservatism in species' traits conferring their environmental tolerances, spatial phylogenetic turnover between early successional communities will (i) be higher than expected given the level of species turnover, (ii) be explained by environmental differences between communities (Bartlett *et al.*, 2015; Cadotte & Tucker, 2017) and iii) be detected only at large phylogenetic scales (Cavender-Bares & Reich, 2012; Hardy *et al.,* 2012). If, in contrast, there is an increase in the relative importance of biotic filtering, due to limiting similarity competition, during succession, we predict that spatial phylogenetic turnover between late successional communities will be (i) less than expected (spatial phylogenetic overdispersion), (ii) detected at small phylogenetic scales, and (iii) unrelated to environmental differences between plots (Bartlett *et al.,* 2015). Alternatively, if hierarchical competition is the predominant force shaping communities during late succession, we predict that late successional communities will be comprised of closely related species, but that phylogenetic turnover will not covary with environmental differentiation (Bartlett *et al.,* 2015). If traits conferring competitive dominance are phylogenetically conserved, and competitively superior species belong to a particular clade (Roeder *et al.,* 2015), we additionally predict that hierarchical competition will cause phylogenetic clustering at shallow phylogenetic scales. In contrast, if late successional communities are primarily governed by the accumulation of closely related species that share adaptations to the local abiotic conditions (Li *et al.,* 2015) and environmental filtering selects for distinct sets of closely related species in plots that differ in their abiotic environment, we predict that spatial phylogenetic turnover between communities belonging to the late successional stages will be (i) higher than expected, (ii) explained by environmental differences between sites, and (iii) detected at broad phylogenetic scales, resulting from phylogenetic conservatism of abiotic niches. Finally, if deterministic community assembly results in temporal shifts in phylogenetic community composition due to successional changes in abiotic conditions (Swenson *et al.,* 2012), we predict that phylogenetic turnover between stages will (i) be higher than expected by chance, (ii) be higher than spatial turnover between plots from the same stage, and (iii) increase with environmental differences between stages. Conversely, if relatively constant abiotic conditions cause a lack of phylogenetic shifts to over time (Letten *et al.,* 2014), we predict that phylogenetic turnover between successional stages will be (i) low relative to species turnover, (ii) lower than phylogenetic turnover between plots from the same stage, and (iii) unrelated to environmental differences between stages.

To test these predictions, we use data on tree communities representing different stages of a subtropical forest succession in south-eastern China. Successional subtropical forests provide an ideal system for the study of temporal changes in the mechanisms underlying community assembly as they represent community assembly in action and are exceptionally species-rich (Uriarte *et al.,* 2010; Arroyo-Rodríguez *et al.,* 2017). While subtropical forest areas were once widespread across South and East China, they are currently under severe decline as a result of land use intensification (Wang *et al.,* 2007). Because of frequent anthropogenic disturbance events, such as logging and burning, subtropical forests often consist of a mosaic of different stages of secondary forest succession. Combining analysis of spatial and temporal turnover (at the individual- and species-level), while examining turnover (i) at different phylogenetic depths and (ii) with increasing environmental differentiation, we will be able to address competing predictions about the temporal changes in the relative importance of the processes that generate biodiversity after disturbance.

## Materials and methods

### Study area and sampling

We studied woody plant communities in the comparative study plots that had been established within the biodiversity-ecosystem functioning experiment BEF-China (Bruelheide *et al.,* 2011). The plots represent a chronosequence of subtropical forest succession in the Gutianshan National Nature Reserve (GNNR), located in Zhejiang Province in south-eastern China (29°8'18"-29°17'29"N, 118°2'14"–118°11'12" E). The GNNR comprises mixed broad-leaved forests (Wu & others, 1980; Hu & Yu, 2008) within an elevational range of 250 m to 1258 m a.s.l‥ A total of 1426 seed plant species of 648 genera and 149 families has been recorded in GNNR (Lou & Li, 1988). The study area mainly consists of a mosaic of secondary forest stands that represent different successional stages, with maximum tree age of approximately 180 yrs (Bruelheide *et al.,* 2011).

Species abundance data was obtained from a vegetation inventory (May-October 2008) of all individuals of trees and shrubs (> 1 m height, 147 species in total) in each of the 27 30×30m plots (see Bruelheide *et al.,* 2011). The plots were distributed over the GNNR to represent five successional stages (differing by 20 years), based on estimations of the age of the largest tree individuals and on knowledge of the last logging event [see Bruelheide *et al. (*2011) for more detailed information on type of disturbance that preceded succession]. The number of plots per successional stage were 5 (< 20 yr), 4 (20-39 yr), 5 (40-59 yr), 6 (60-79 yr) and 7 (≥ 80 yr). Because fewer individuals were recorded in the older plots relative to the younger plots (Fig. S1 in Supporting Information), we assessed whether the differences in the number of individuals between plots may potentially bias our results, which was not the case in our study (Table S1).

For each plot, a set of environmental variables (Table S2) related to topography [aspect (expressed as northness and eastness), slope, elevation], light (photosynthetically active radiation (PAR), red/far-red ratio) and soil characteristics (pH, moisture, C/N-ratio) were available from Bruelheide *et al.* (2011) and Kröber *et al.* (2012). Total phosphorus (P) content of the soil was measured with nitric acid digestion, a standard method recommended by the German forest soil survey (BMELV, 2009). The inorganic nitrogen concentration (NO3^-^, NH4^+^) of the mineral soil was determined by KCl extraction (1mol/L) followed by Flow Injection Analysis (FIAstar 500 Analyzer, FOSS, Hilerød, Denmark).

### Phylogenetic data and regional species pool

Based on the set of species present in the 27 plots and on the list of all woody species of the Gutianshan National Nature Reserve (Lou & Li, 1988), we constructed a regional species pool [the set of 438 woody species that occur in the whole GNNR (Table S3)] for which a phylogeny was inferred. For details on phylogenetic inference see Methods S1 and Tables S4 & S5. In short, we obtained sequence information (matK, rbcL and ITS region) for all species, or their closest relatives, from GenBank or de novo using standard barcoding protocols. A maximum likelihood tree was computed and dated using non-parametric rate smoothing and using published fossils as age constraints (Methods S2, S3). To avoid potential bias in the analysis of phylogenetic patterns due to their disproportionately long branch lengths (Letcher, 2010; Cadotte, 2014), non-angiosperm and one bamboo (*Pleioblastus amarus,* Poaceae) species, which generally occurred at low frequencies within the study area, were excluded from the regional species pool. We further excluded cultivated species, resulting in a total of 410 woody species of which 143 occurred in the 27 study plots (Table S3).

### Phylogenetic structure

Using information on species composition and the phylogenetic tree pruned down to the 143 woody angiosperms found in the 27 plots, we estimated phylogenetic structure following the framework proposed by Hardy & Senterre (2007), which is based on the spatial decomposition of evolutionary relatedness between species into within- and between-community components. Within the Hardy & Senterre (2007) framework, spatial phylogenetic structure was quantified for presence/absence and abundance data, using the phylogenetic turnover (between-plot differentiation) statistics П_ST_ and BST, respectively: [inline] and [inline] where 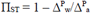 and 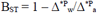 alpha diversity, and correspond to the mean within-community phylogenetic distance between distinct species and the mean phylogenetic distance between two individuals of distinct species, respectively, averaged over all communities belonging to the same successional stage. 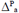 and 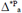 are the mean phylogenetic distance between distinct species and the mean phylogenetic distance between two individuals of distinct species, respectively, sampled from different communities belonging to a particular stage. Values of spatial phylogenetic turnover, П_ST_ or B_ST_, > 0 indicate spatial phylogenetic clustering – species, or individuals, within communities are phylogenetically more related than species, or individuals, from different communities. Spatial phylogenetic overdispersion is observed if П_ST_ or B_ST_ < 0, indicating that species, or individuals, within communities are phylogenetically less related than species, or individuals, from different communities. When П_ST_ and B_ST_ are calculated between pairs of plots belonging to the same successional stage, they address within-stage phylogenetic turnover. When П_ST_ and B_ST_ are calculated between pairs of plots belonging to different successional stages, they address between-stage phylogenetic turnover. We tested, based on 100 simulation runs, whether levels of spatial phylogenetic turnover were affected by differences in the number of plots among stages (Methods S4). Mean Pearson correlations between П_ST_ (or B_ST_) for simulated communities and the number plots were close to zero, indicating that levels of phylogenetic turnover were not simply a reflection of the number of plots. To complement our main analyses of phylogenetic turnover, and in addition to measures of phylogenetic alpha diversity (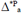 and 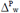), we also calculated Shannon evenness (Magurran, 2004) for each plot.

### Null models

To test whether П_ST_ or B_ST_ were significantly higher (or less) than zero, observed П_ST_ or B_ST_ values were compared to those re-calculated for 999 random communities. Random communities were generated using null model '1p' in Hardy (2008), shuffling species names across the phylogeny of all 410 woody angiosperms from the regional species pool. The latter corresponding to the set of species that are present in, or could potentially colonize, our study plots (see Ding *et al.,* 2012 and Letcher *et al.,* 2012). This null model maintains (i) the number of species within each community, (ii) species turnover between communities, (iii) the patterns of spatial autocorrelation in overall species abundances and occurrence frequencies, (iv) species' occurrence frequency across the study landscape and (v) species identity within each successional time step. This type of null model is appropriate for temporal data (Letcher *et al.,* 2012; Norden *et al.,* 2012) and has been demonstrated to provide exact tests (i.e. correct Type-I error rates) in situations where overall species frequencies (or abundances) are not phylogenetically structured (Hardy, 2008, see Methods S5). Significant positive (or negative) values of П_ST_ (or B_ST_) of within-stage phylogenetic turnover indicate that species, or individuals, co-occurring within successional stages are more (or less) related than expected by chance. Higher-than-expected П_ST_- or B_ST_-values of between-stage phylogenetic turnover that are higher than within-stage phylogenetic turnover indicate phylogenetic shifts during the course of succession. Lower-than-expected values of between-stage turnover, that are lower than within-stage turnover, would indicate constant phylogenetic composition during succession.

### Phylogenetic structure at different depths in the phylogeny

We assessed whether non-random phylogenetic structure, within each of the five successional stages, occurred at particular phylogenetic depths, following the approach in Hardy & Senterre (2007): phylogenetic turnover between plots was calculated based only on species pairs within clades younger than a given divergence time threshold. We chose eleven age thresholds, ranging between 30 Myr to 128 Myr, by steps of approximately 10 Myr. To test whether phylogenetic turnover significantly differed from zero at particular phylogenetic scales, we carried out partial randomizations, shuffling species names across the phylogeny, but restricting the randomization to species within clades younger than the respective age threshold. All calculations of phylogenetic community structure were carried out on phylogenetic, cophenetic distance matrices, using the packages 'vegan' (Oksanen *et al.,* 2017) and 'spacodiR' (Eastman *et al.,* 2011) in the R statistical package (R Development Core Team, 2017) and SPACoDi 0.10 (Hardy, 2010). To identify clades that signifcantly contributed to phylogenetic turnover between plots, we tested for each node in the phylogeny whether it had more decendent taxa than expected in a particular plot, using the 'nodesig' procedure in Phylocom v.4.2 (Webb *et al.,* 2009).

### Relating phylogenetic structure to environmental variables

To quantify the extent to which spatial and temporal phylogenetic turnover was explained by differences in abiotic conditions, pairwise П_ST_ (or B_ST_) values were regressed on between-plot environmental distances. To control for covariation between phylogenetic turnover and spatial distance, we used the residuals from regressions of П_ST_ (or B_ST_) against the Euclidean distances calculated from the geographic x- and y-coordinates of the plots instead of the actual phylogenetic turnover values. Significance of the relationships was assessed by non-parametric randomization testing [5000 randomizations, R-package 'lmPerm' (Wheeler & Torchiano, 2016)]. Environmental distances were obtained from an inter-plot distance matrix based on the 11 topographic, light and edaphic descriptors. A principal components analysis (PCA) was carried out on the log-transformed and standardized (mean = 0, sd = 1) environmental data, to correct for the dominance of the distance matrix by highly correlated environmental variables. The resulting first six principal components (PCs) accounted for about 90% of the total variation (Table S6) and were used to construct the Euclidean inter-plot distance matrix from which the environmental distances were obtained. Because associations between phylogenetic turnover and environmental differentiation may be a reflection of differences in sample size among the successional stages, we additionally assessed relationships between environmental and phylogenetic turnover at each stage based resampling all possible combinations of four plots, the minimum number of plots across stages.

### Phylogenetic signal in traits

To assess whether phylogenetic relatedness between species reflects their ecological similarity, we quantified phylogenetic signal in six traits [leaf area, specific leaf area (SLA), leaf nitrogen content, leaf phosphorus content, wood density, maximum height] that represent multiple axes of plant functional differentiation (Westoby *et al.,* 2002; Wright *et al.,* 2004; Chave *et al.,* 2009; Moles *et al.,* 2009). Estimates of phylogenetic signal were based on the three metrics Blomberg’ *K* (Blomberg *et al.,* 2003), Pagel's *λ* (Pagel, 1999) and Abouheif/Moran's *I* (Abouheif, 1999) (Table S7), and calculated in the R-packages 'phytools' (Revell, 2012) and 'adephylo' (Jombart *et al.,* 2010), for the subset of 121 species (of the 143 angiosperm species occurring in the 27 plots) for which data on all six traits were available from Kröber *et al.* (2012) and Böhnke *et al.* (2012, 2014).

## Results

### Temporal changes in alpha diversity

Phylogenetic alpha diversity (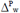 and 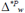) showed no significant temporal trend in the course of succession (Fig. 1a,b). In contrast, there was a steep increase in species (Shannon) evenness over time (Fig. S2).

**Fig. 1.**
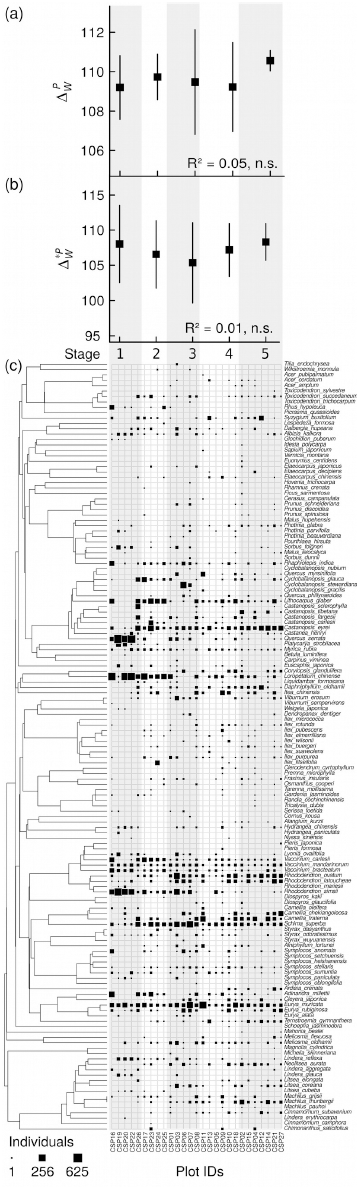
Phylogenetic alpha diversity within the five successional stages (mean ± 1 SE; Stage 1 (< 20 yr): n=5, Stage 2 (20-39 yr): n=4, Stage 3 (40-59 yr): n=5, Stage 4 (60-79 yr): n=6, Stage 5 (≥ 80 yr): n=7), based on (a) presence/absence (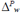) and (b) abundance data (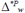). 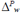 and 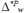 are equivalent to the mean phylogenetic distance between distinct species (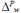)and the mean phylogenetic distance between individuals of distinct species 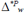 within communities, respectively. R^*2*^ values are given. None of the two alpha diversity measures showed a significant successional trend. (c) Distribution of abundances within the 27 comparative study plots [assigned to one of the five successional stages (Stage 1-5)] mapped onto the phylogeny of the 143 species. The size of the black squares corresponds to the number of individuals.

### Comparisons between spatial and temporal phylogenetic turnover

Levels of overall phylogenetic turnover were significantly different from those predicted, given the levels of species turnover (Fig. 2). However, deviation from null expectations showed opposing patterns depending on whether phylogenetic turnover was estimated based on species presence/absence (П_ST_) or abundance (B_ST_). Overall levels of presence/absence-based turnover were higher than expected, whereas overall abundance-based turnover was lower than expected. When overall phylogenetic turnover was dissected into turnover between pairs of plots belonging to the same successional stage (within-stage spatial turnover) and turnover between pairs of plots at different successional stages (between-stage temporal turnover) respectively, presence/absence-based within-stage turnover (П_ST_) was higher than expected, indicating that species within plots were more closely related to each other than to species from different plots. Levels of presence/absence-based between-stage turnover (_ПST)_ did not differ from random expectations (Fig. 2a). In contrast, between-stage turnover was on average lower than predicted by chance, when based on abundance data (B_ST_).

**Fig. 2.**
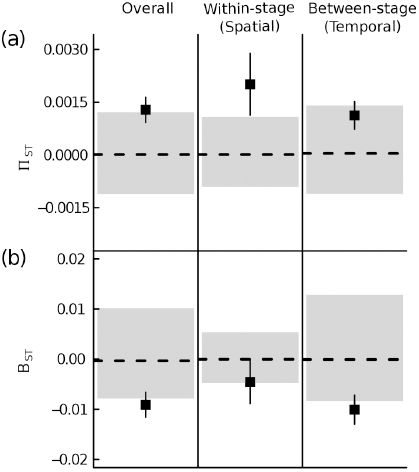
Phylogenetic turnover for all pairs of plots (combining spatial and temporal turnover, n=351, left panel) dissected into spatial, i.e. within-successional stage, (n=62, middle panel) and temporal, i.e. between-stage, (n=289, right panel) turnover (black squares, mean ± 1 SE). Phylogenetic turnover was calculated for (a) presence/absence (П_ST_) and (b) abundance data (B_ST_) and is based on the partitioning of the mean phylogenetic distance between distinct species, or between individuals of distinct species, into within- and between-community components. П_ST_ or B_ST_ > 0 indicate that the species, or individuals, co-occurring within communities are phylogenetically more related to each other than to species from other communities (high turnover). B_ST_ or П_ST_ < 0 indicate that the species, or individuals, co-occurring within communities are phylogenetically less related to each other than to species from other communities (low turnover). The black dashed line and grey-shaded area represent the mean and the 95% CI, respectively, from the 999 random communities. B_ST_ and П_ST_ values outside the interval indicate non-random phylogenetic turnover.

### Phylogenetic turnover within and between single successional stages

Spatial phylogenetic turnover measures showed contrasting patterns of deviation from random expectations over the course of succession (Fig. 3). Presence/absence-based phylogenetic turnover (П_ST_) did not significantly differ from zero within early and mid successional stages (stages 1, 2 and 3, Fig. 3a). However, П_ST_-values were higher than expected within the two latest successional stages (stages 4 and 5, Fig. 3a). In contrast, abundance-based spatial phylogenetic turnover (B_ST_) was lower than predicted by chance within the first successional stage but did not significantly differ from null expectations within the mid-and late-successional stages (Fig. 3b). Presence/absence-based turnover (П_ST_) between pairs of consecutive successional stages was higher than expected between the mid and last successional stages (stage 3-4, stage 3-5 and stage 4-5, Fig. S3), but was never higher than levels of turnover within each of the stages 3, 4 and 5 (Fig. 3a). Presence/absence-based turnover (B_ST_) was lower than predicted between the early and mid successional stage as well as between the first and the last stage (stage 1-2 and stage 1-5, Fig. S3), with values of B_ST_ that were lower than those estimated within stages (Fig. 3b).

**Fig. 3.**
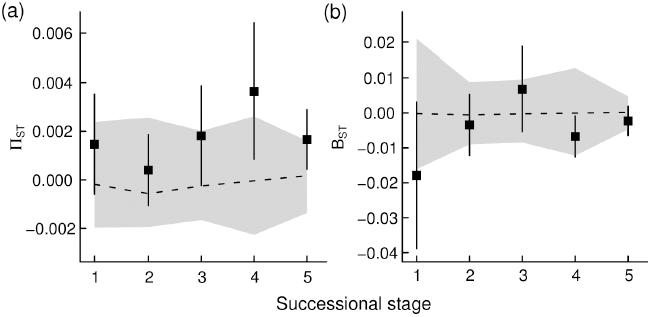
Spatial phylogenetic turnover between all pairs of communities within each of the five successional stages (black squares, mean ± 1 SE; Stage 1: n=10, Stage 2: n=6, Stage 3: n=10, Stage 4: n=15, Stage 5: n=21), based on (a) presence/absence (П_ST_) and (b) abundance data (B_ST_). B_ST_ or П_ST_ values above (or below) the grey-shaded area (i.e. the 95% CI for the BST or ПST values from the 999 random communities) indicate spatial phylogenetic clustering (or overdispersion).

### Covariation between phylogenetic turnover and environmental differentiation

There were no significant relationships of presence/absence-based overall phylogenetic turnover and between-stage phylogenetic turnover (П_ST_), respectively, with environmental differences between plots (Fig. S4a,c). Instead, there was on average a significant positive association between within-stage phylogenetic turnover (П_ST_) and environmental distance (Fig. S4b), indicating an increase in phylogenetic turnover with increasing environmental differences (mainly related to soil moisture and light, see Table S6 & Fig. S7), between plots that belong to the same successional stage. When relationships between П_ST_ and environmental distance were assessed within each of the five successional stages separately, significant increases in phylogenetic turnover with increasing environmental distance were only detected within the two last successional stages (stage 4 and 5, Fig. 4). The significant positive associations between phylogenetic turnover and environmental differences between plots within the two latest successional stages were maintained after accounting for differences in sample size between the stages using resampling down to the minimum number of plots (n=4) across stages (Stage 4: R^2^=0.24^*^; Stage 5: R^2^=0.19^*^). Abundance-based phylogenetic turnover (B_ST_) was not associated with environmental distances, neither within nor between successional stages (results not shown).

**Fig. 4.**
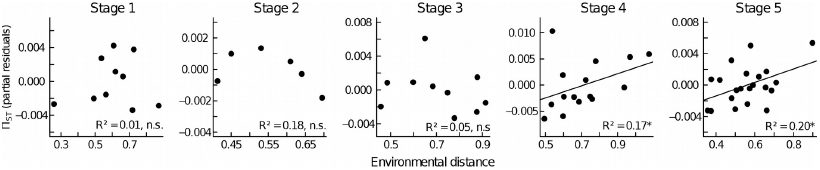
Relationship between presence/absence-based phylogenetic turnover (П_ST_) and environmental differences (with respect to topography, light and soil characteristics) between communities, within each of the five successional stages. П_ST_ values are given as partial residuals after accounting for spatial distance as a covariable. R^2^ values are given. Significant relationships (based on randomization testing) are indicated by solid lines and are only detected in the two late successional stages. ^*^ P ≤ 0.05, n.s. not significant.

### Phylogenetic structure at different depths in the phylogeny

Presence/absence-based phylogenetic turnover (П_ST_) within the early and mid successional stages did not differ from random expectations throughout the phylogeny (Fig. 5). Non-random and higher-than expected phylogenetic turnover was only detected within the two latest successional stages (stage 4 and 5, Fig. 5) and occurred close to the root of the phylogeny (> 100 Myr), indicating phylogenetic clustering at a deep phylogenetic scale. Abundance-based phylogenetic turnover (B_ST_) did not differ from random expectations at any level in the in phylogeny within any successional stage (results not shown). Clades that were over-represented in, and contributed to the high turnover between, pairs of plots within the late successional stages diverged early in phylogeny (~ 100 Myr ago). Nodes that were significantly associated (i.e. had more taxa than expected by chance) with each of the plots are listed in (Table S8). For instance, the plot pair with the highest level of phylogenetic turnover within the late successional stage 4 (plot IDs CSPs 5 and 11), (i) had significantly more taxa than expected within the families Ericaceae (*Rhododendron, Vaccinium, Lyonia, Pieris*) and Theaceae (*Camellia, Schima*) (nodes 44 & 39) that diverged within the Ericales ~ 100 Myrs ago (Fig. S6) and (ii) was associated with dry and moist soil conditions, respectively (Fig. S7).

**Fig. 5.**
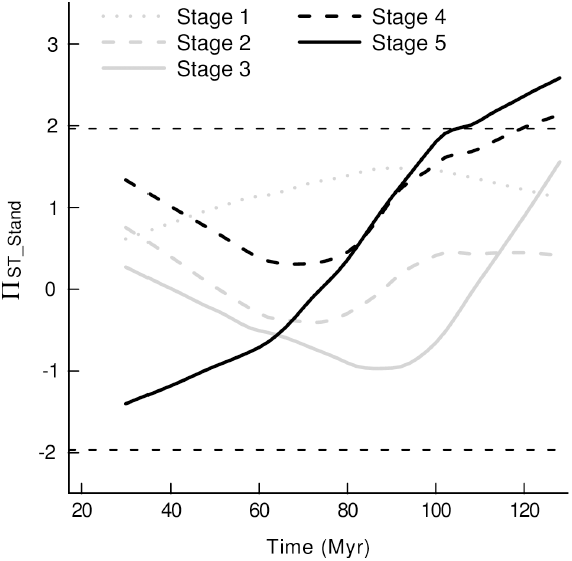
Phylogenetic turnover, based on presence/absence data (П_ST_), at different phylogenetic depths, within the five successional stages. The lines represent, for each successional stage, fitted curves from local polynomial regression (loess, smoothing span = 0.66, polynomial degree = 1), of node age against the standardized effect size of phylogenetic turnover (П_ST_Stand_). П_ST_Stand_ values were calculated as the ratio between observed to expected values of П_ST_: П_ST_Stand_=(П_ST_obs_-П_ST_exp_)/sd(П_ST_exp_), where П_ST_obs_ is the observed П_ST_ value at a particular node, and П_ST_exp_ and sd(П_ST_exp_) are the mean and standard deviation of the expected П_ST_ values from 999 partial phylogenetic tree randomizations among clades younger than that particular node. The two horizontal dashed lines indicate the 0.05 significance levels. Non-random and higher-than-expected turnover (spatial phylogenetic clustering) was only detected within the two late successional stages and at broad phylogenetic scales (from approximately 128 to 100 Myr).

### Phylogenetic signal in traits

All of the six traits considered showed significant phylogenetic signal, with values of Blomberg's *K*, Pagel's *λ* and Abouheif/Moran's *I* significantly greater than expected from a null model of no phylogenetic signal (Table S7). This suggests that, in our study, phylogenetic relatedness reflects overall trait similarity.

## Discussion

The present study combines analysis of within- and between-stage phylogenetic turnover during succession across phylogenetic scales, while accounting for between-plot environmental differentiation, and demonstrates that, despite a lack of temporal phylogenetic turnover between stages, there was a shift from abundance-based phylogenetic overdispersion in early succession towards presence/absence-based phylogenetic clustering in late succession. Low between-stage turnover that was not explained by environmental differences between stages suggests that (i) relatively constant environmental conditions and (ii) shifts in species abundances (towards higher evenness) that were counterbalanced by increasing relatedness towards late succession, resulted in an absence of net change in phylogenetic composition over time. Within the late successional stages, phylogenetic turnover was higher than expected, increased with environmental differentiation between sites and occurred at broad phylogenetic scales, indicating (i) deep phylogenetic conservatism of species' abiotic niches, and (ii) that environmental filtering along an abiotic gradient becomes more important towards late succession.

### Comparisons between spatial and temporal phylogenetic turnover: high turnover within and low turnover between successional stages

Within-stage and between-stage phylogenetic turnover showed, on average, opposing levels of deviation from random, depending whether they were based on presence/absence or abundance data. While turnover between plots belonging to the same successional stage was higher than expected, relative to the levels of species turnover, when based on presence/absence data, phylogenetic turnover between plots at different successional stages was lower than expected when based on abundance data (Fig. 2). Preceding studies (Lozupone *et al.,* 2007; Fine & Kembel, 2011) have demonstrated that using both presence/absence- and abundance-based metrics may reveal different patterns of phylogenetic structure for rare and abundant species, and thus may help to distinguish species composition from dominance effects. The previous study by Norden *et al.* (2012) revealed that temporal changes in phylogenetic community structure during tropical rainforest succession were influenced by shifts in species' abundance rather than species occurrence, whereas Letten *et al.* (2014) found low temporal phylogenetic turnover during heathland succession, because closely related, dominant species replaced each other over time. The previous study of Bruelheide *et al.* (2011), in the same system that was used in our study, demonstrated a lack of species turnover with only few species restricted to a particular successional stage, reminescent of the concept of initial floristic composition, but that there were substantial shifts in species' abundance towards a more even distribution of abundance in late successional communities. Therefore, in our study, the low levels of abundance-based phylogenetic turnover, relative to the turnover of species between successional stages, reflect the fact that the temporal increase in evenness is counterbalanced by the increase in relatedness between the most dominant species towards late succession (Figs. 1c & S2): the most dominant species within the early successional stages *(Loropetalum chinense, Quercus serrata, Rhododendron simsii*) are distantly related, whereas late successional communities were comprised of closely related species, i.e. belonging to the genera *Castanopsis, Rhododendron, Camellia* and *Eurya*, respectively – resulting in an absence of a net change in phylogenetic diversity and composition over time. Further, low levels of temporal functional turnover during tropical forest succession, were detected in an earlier study by Swenson *et al.* (2012), presumably due to relatively constant local environmental conditions through time. In our study, environmental differences between communities at different successional stages were similar to those between communities at the same stage (Fig. S4b,c), indicating that the lack of phylogenetic shifts likely reflects the constant abiotic conditions throughout succession.

In spite of the lack of temporal phylogenetic turnover between stages, we found a higher than expected presence/absence-based phylogenetic turnover (П_ST_) between plots that belong to the same successional stage, suggesting that there are filtering processes that have selected for different groups of closely related species. Our finding that the within-stage phylogenetic turnover (П_ST_) significantly increased with environmental distance (Fig. S4b) indicates that phylogenetic differentation between communities belonging to the same successional stage was due to an underlying environmental gradient (mainly related to soil moisture and light; see Table S6), and that the higher-than-expected levels of spatial phylogenetic turnover reflect differential abiotic filtering selecting for closely related species within communties that belong to the same successional stage (see following section). The strong association between within-stage phylogenetic turnover and environmental differences may also be a reflection of the fact that, in contrast to previous studies of community turnover in subtropical forest systems that have focussed on indirect abiotic descriptors such as elevation or habitat types (Legendre *et al.,* 2009), we used a large set of environmental (edaphic, light & topographic) descriptors. And it has been demonstrated recently that the quality of environmental data may influence conclusions about assembly processes (Chang *et al.,* 2013).

### Temporal changes in within-stage turnover

We found that there was a shift from (abundance-based) spatial phylogenetic overdispersion within the first successional stage towards (presence/absence-based) spatial phylogenetic clustering within the two late successional stages (Fig. 3). This contrasts with a number of previous studies of successional tropical and subtropical forests (Letcher, 2010; Ding *et al.,* 2012; Norden *et al.,* 2012; Whitfeld *et al.,* 2012) that found high levels of phylogenetic relatedness in young, disturbed forest communities, compared to older communities. Those studies concluded that disturbance in early succession acts as an abiotic filter and selects for closely related species but that competitive exclusion of closely related species becomes increasingly important towards late succession. Our finding that the most dominant species within plots were less related to each other than to species from different plots within the first successional stage may be explained in a number of different ways: First, phylogenetic overdispersion may reflect abiotic filtering if the traits conferring environmental tolerance are not phylogenetically conserved and distantly related species are filtered by the same environment (Cavender-Bares *et al.,* 2004). However, we detected significant phylogenetic signal in a set of six traits reflecting multiple axes of plant functional differentiation, and Eichenberg *et al.* (2015) found even stronger phylogenetic signal in the same study system when intraspecific trait variation was taken into account. This indicates that phylogenetic relatedness reflects ecological similarity between species and that abiotic filtering of convergent niche traits is unlikely to explain phylogenetic overdispersion in our study. Second, phylogenetic overdispersion may result from competitive exclusion of closely related species that share similar traits – a process that is expected to result in overdispersion at small phylogenetic scales. However, in our study, we did not detect phylogenetic overdispersion at shallow phylogenetic depth (Fig. 5). Third, it has recently been demonstrated that early-successional communities may be comprised of distantly related species in cases where (i) early-successional pioneers are distributed all over the phylogeny (Letcher *et al.,* 2015) and/or (ii) remnant species, which have persisted from former management, have a wide range of phylogenetically conserved traits that allow them to tolerate early successional environmental conditions (Bhaskar *et al.,* 2014). Because in our study, (i) most species were present throughout succession, and (ii) remnant species were represented by only a few individuals (e.g. *Nyssa sinensis, Castanea henryi, Cyclobalanopsis glauca, Castanopsis fargesii*; see Fig. 1c & Bruelheide *et al.,* 2011) and hence did not substantially contribute to abundance-based phylogenetic structure, the abundance-based phylogenetic overdispersion in early succession is unlikely to reflect the presence of pioneer or remnant species. Finally, phylogenetic overdispersion may reflect successful dispersal of species that have different dispersal stategies (Du *et al.,* 2012), provided that dispersal traits are phylogenetically conserved (Baeten *et al.,* 2015). In our study, the most abundant species within the first successional stage (e.g. *Loropetalum chinense, Quercus serrata, Rhododendron simsii*) were both, distantly related (Fig. 1c) and dispersed by different dispersal modes (animal-dispersed acorns, ballistic- and wind-dispersed seeds for *Quercus, Loropetalum* and *Rhododendron*, respectively), suggesting that the abundance-based phylogenetic overdispersion in early succession likely reflects the coexistence of a wide range of different dispersal strategies (Levin & Muller-Landau, 2000; Purschke *et al.,* 2014).

Within the two late successional stages, presence/absence-based phylogenetic turnover was higher than expected relative to the levels of species turnover, indicating deterministic filtering that selects for distinct sets of closely related species in the different plots. There are a few studies that found increasing functional similarity in (sub-)tropical forest communities over time (Uriarte *et al.,* 2010; Buzzard *et al.*, 2015), concluding that the relative importance of abiotic filtering increases with forest age. Further, the previous studies by Hardy *et al.* (2012) and Fine & Kembel (2011), focussing on phylogenetic turnover between tree communities along environmental gradients, pointed out that, if environmental niches are evolutionarily conserved, abiotic filtering is predicted to result a in strong covariation between phylogenetic turnover and environment differentiation between plots (Cadotte & Tucker, 2017). Therefore, our finding that phylogenetic turnover within late successsional stages was higher than expected and explained by environmental differentation [mainly related to to soil and light conditions (Table S6), and independent of spatial distance] between plots (Fig. 4), is consistent with phylogenetic niche conservatism and indicates that the relative importance of environmental filtering along an environmental gradient increased during the course of succession. The high phylogenetic turnover within the late successional stages, together with the lack of temporal between-stage phylogenetic turnover, further suggests that phylogenetic clustering in late succession reflects the local colonization of species that (i) are closely related to residents (Li *et al.,* 2015) and (ii) were already present in the early-successional species pool, indicating that species sorting into their preferred habitat takes time to develop. Spatial phylogenetic clustering in late succession was only detected close to the root of the phylogenetic tree (Fig. 5). Previous studies of community turnover across phylogenetic scales (Parmentier & Hardy, 2009; Cavender-Bares & Reich, 2012) found that phylogenetic turnover increased both with phylogenetic depth as well as with environmental differentiation between sites, and concluded that ancient diversification events, together with niche conservatism, still show an imprint on the assembly of current plant communities. The fact that, in our study, spatial phylogenetic clustering (П_ST_) within late successional communities was only detected at large phylogenetic scales (i.e. between taxa that diverged > 100 Myrs years ago), together with the finding that phylogenetic turnover was explained by abiotic differences (related to soil and light conditions) between plots is consistent with deep phylogenetic signal in species' soil moisture and light niche. Clades that contributed to the high phylogenetic turnover within the late successional stage diverged early in phylogeny and were associated with one or the other end of the environmental gradient (Table S8, Fig. S7), indicating environmental niche differentiation between species that diverged early in phylogeny.

Alternatively, phylogenetic clustering in late succession can also result from hierarchical competition if early successional pioneers are replaced by competitively superior closely related species in late succession (Kunstler *et al.,* 2012; Letten *et al.,* 2014). However, most early-successional species in our study were still present in late succession. Further, hierarchical competition is predicted to result in phylogenetic clustering that is unrelated to environmental differentiation between plots (Bartlett *et al.,* 2015), which was not the case in our study. This suggests that competition hierarchies are unlikely to explain the phylogenetic clustering in our study. Our finding that non-random phylogenetic structure within the two latest successional stages was only detected based on presence/absence-data (Fig. 3a), is likely to reflect the high number of rare species found in late compared to early succession (Fig. 1c, see also Bruelheide *et al.,* 2011), and in such situations presence/absence metrics (such as П_ST_), giving high weight to rare species, will provide greater testing power to detect significant community phylogenetic structure than metrics based on abundance (Helmus *et al.,* 2007; Vellend *et al.,* 2011).

In conclusion, the integrated analysis of the spatial and temporal components of phylogenetic relatedness during succession, across phylogenetic and environmental scales, allowed to test competing hypothesis about the temporal dynamics of community processes after disturbance. Our results do not support a model that predicts a progression towards decreasing phylogenetic relatedness over time. Instead, our findings support a deterministic model of community assembly where the phylogenetic composition is constrained though time but different assembly processes act at different ends of the successional gradient: colonization of species that differ in their dispersal strategies likely plays an important role in early succession, whereas, despite the lack of phylogenetic shifts between stages, environmental filtering of niche traits that are conserved deep in phylogeny becomes increasingly important towards late succession. Such insights into the temporal dynamics of post-disturbance community assembly processes were not apparent from previous analyses that focused either on single (spatial or temporal) phylogenetic turnover components or single phylogenetic scales.

## Acknowledgements

We would like to thank the administration and the staff (in particular Fang Teng for species identification and abundance assessment for the species pool) of the Gutianshan National Nature Reserve, and the members of the BEF China consortium, for their support, Stefan Trogisch, Michael Scherer-Lorenzen, Björn Todt and Jü rgen Bauhus for providing data on soil chemistry, Andreas Prinzing, Nathan G. Swenson and Antoní n Machá č for discussions and comments on an earlier version of the manuscript. The study was financed by the German Research Foundation (DFG FOR 891/1-3 and BR 1698/9-1-3). O.P. acknowledges the support by the German Centre for Integrative Biodiversity Research (iDiv) Halle-Jena-Leipzig, funded by the German Research Foundation (FZT 118).

## Author contributions

H.B. established the BEF-China experimental platform. O.P. developed the main idea for this manuscript with contributions from W.D., S.G.M. & H.B. O.P. analysed the data and interpreted the results with input from all co-authors. S.G.M. generated the phylogenetic tree with contributions from W.D. O.P. wrote the first draft of the manuscript with all other authors substantially contributing to revisions.

## Supporting Information

**Fig. S1.**
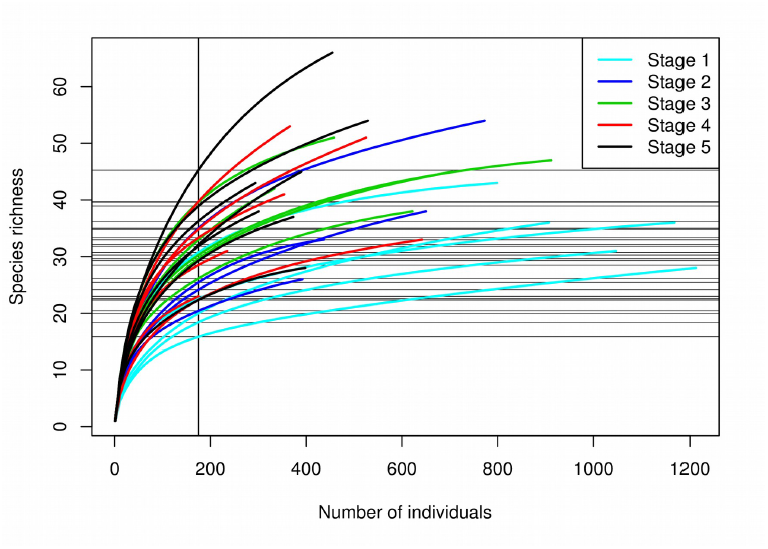
Rarefaction curves of the 27 woody plant communities (CSPs), giving the estimated number of species for any number of individuals. The five successional stages are indicated by different line colors. The vertical line depicts the minimal number of individuals (n=175) sampled in a plot. The intersection between the rarefaction curves and the vertical line corresponds to the estimated number of species if only 175 individuals per plot were sampled.

**Fig. S2.**
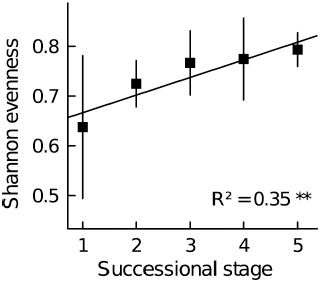
Shannon evenness within each of the five successional stages (black squares, mean ± 1 SE). R^2^-value is given. The solid line indicates the significant relationship between evenness and successional stage. ^**^ P ≤ 0.01.

**Fig. S3.**
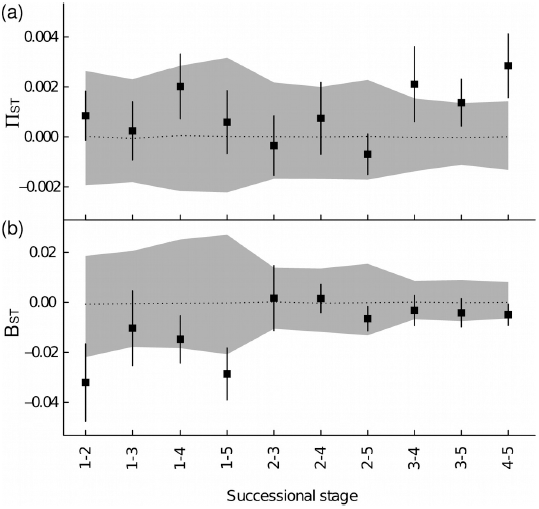
Phylogenetic turnover between successional stages (black squares, mean ± 1 SE), calculated for (a) presence/absence (П_ST_) and (b) abundance data (B_ST_). The black dashed line and grey-shaded area represent the mean and the 95% CI, respectively, from the 999 random communities. B_ST_ and П_ST_ values above the interval indicate higher than expected temporal phylogenetic turnover. BST and П_ST_ values below the interval indicate lower than expected temporal phylogenetic turnover.

**Fig. S4.**
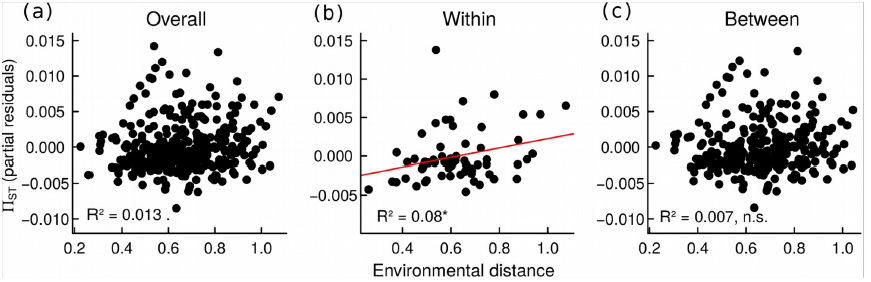
Relationships between presence/absence-based phylogenetic turnover and environmental differences (with respect to topography, light and soil characteristics) between communities for (a) all pairs of plots (combining spatial and temporal turnover, n=351), (b) pairs of plots of the same successional stage (spatial turnover, n=62) and (c) pairs of plots belonging to different successional stages (temporal turnover, n=289). П_ST_ values are given as partial residuals after accounting for spatial distance as a covariable. R2 values are given. The significant relationship (based on randomization testing) between spatial phylogenetic turnover and environmental distance is indicated by the solid red line. ^*^ P ≤ 0.01,. P ≤ 0.1, n.s. not significant.

**Fig. S5.**
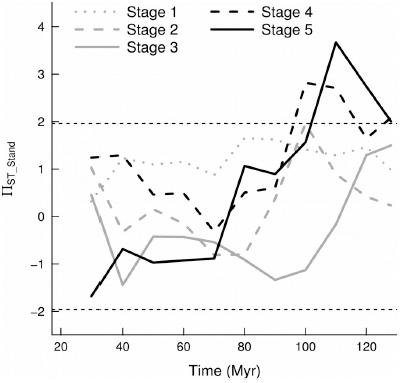
Phylogenetic turnover, based on presence/absence data (П_ST_), at different phylogenetic depths, within the five successional stages. Standardized П_ST_ values (П_ST_Stand_) are given, calculated as the ratio between observed to expected values of ПST: П_ST_Stand_=(П_S_T__obs_-П_ST_exp_)/sd(П _ST_exp_), where П_ST__obs is the observed П_ST_ value at a particular node, and П_ST_exp_ and sd(П_ST_exp_) are the mean and standard deviation of the expected П_ST_ values from 999 partial phylogenetic tree randomizations among clades younger than that particular node. The dashed lines indicate the 0.05 significance levels. Non-random and higher-than-expected turnover (spatial phylogenetic clustering) was only detected within the two late successional stages and at broad phylogenetic scales (from approximately 128 to 100 Myr).

**Fig. S6.**
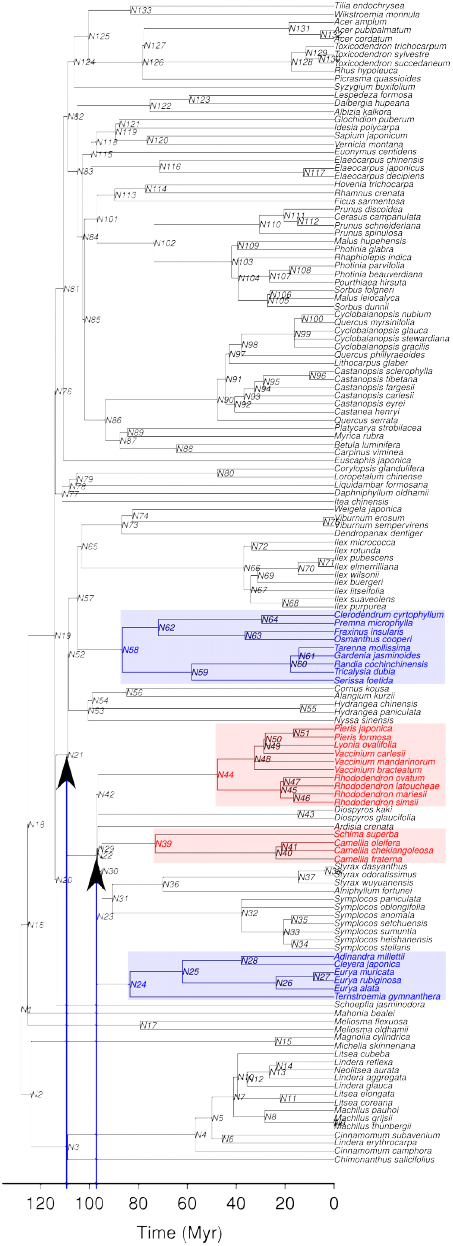
Illustration of results from the nodesig analysis. Highlighted are clades (shaded areas) that had significantly more taxa than expected in plot pairs with the highest levels of phylogenetic turnover at the two latest successional stages (blue: plot pair CSP 5 & 11 at stage 4; red: CSPs 4 & 12 at stage 5, see also Fig. 1c). For instance, node N39 and N44, respectively, i) were significantly associated with the plots CSPs 5 and 11 (the plot pair that that had the highest phylogenetic turnover in stage 4, see also Table S8), and ii) correspond to the families Theaceae (*Camellia, Schima*) and Ericaceae (*Rhododendron, Vaccinium, Lyonia, Pieris*), that diverged early in phylogeny ~ 100 Myrs ago within the Ericales at node N22 (red vertical line). See Table S8 for a complete list of nodes that were significantly associated with each of the plots.

**Fig. S7.**
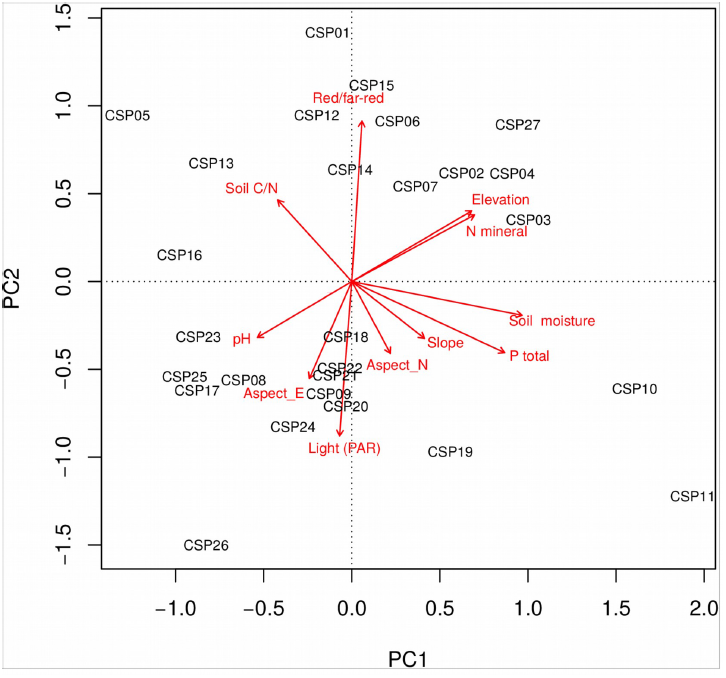
PCA biplot illustrating the association between the 11 environmental variables and the 27 plots (CSPs). See Table S6 for variable loadings and Table S2 for Pearson correlations.

**Table S1.** Correlations between non-rarefied and rarefied phylogenetic metrics. Estimates of phylogenetic diversity and turnover were recalculated (100 times) for rarefied communities containing 175 individuals each (the minimum number of individuals recorded in a plot). Rarefied and non-rarefied estimates for all of the metrics were strongly (P < 0.05) correlated.

| Correlation |  |  |
| --- | --- | --- |
| Metric | Mean | SD |
| $\Delta^P_w$ | 0.821 | 0.036 |
| $\Delta^{*P}_w$ | 0.970 | 0.006 |
| $\Pi_{ST}$ | 0.632 | 0.049 |
| $B_{ST}$ | 0.967 | 0.004 |

**Table S2.** Pearson correlations between successional stage and the 11 abiotic environmental descriptors and successional stage. Significant correlations (P < 0.05) are highlighted in bold.

|  |  |  |  |  | Light | Red/far- | Soil |  |  | N |  |
| --- | --- | --- | --- | --- | --- | --- | --- | --- | --- | --- | --- |
|  | Elevation | Aspect E | Aspect N | Slope | (PAR) | red | moisture | pH | Soil C/N | mineral | P total |
| Stage | 0.29 | -0.26 | -0.26 | 0.11 | **-0.41** | **0.59** | 0.29 | -0.36 | 0.1 | **0.59** | 0.07 |
| Elevation |  | -0.29 | 0.03 | -0.11 | -0.28 | 0.21 | **0.51** | **-0.43** | -0.06 | **0.47** | 0.29 |
| Aspect_E |  |  | 0.24 | 0.23 | **0.39** | -0.31 | -0.21 | 0.11 | 0.1 | -0.18 | 0.03 |
| Aspect_N |  |  |  | **0.4** | 0.21 | -0.21 | 0.15 | -0.26 | 0.13 | -0.18 | 0.28 |
| Slope |  |  |  |  | 0.04 | -0.06 | 0.33 | -0.15 | 0.05 | 0.14 | **0.48** |
| Light (PAR) |  |  |  |  |  | **-0.82** | 0.14 | 0.09 | -0.27 | -0.25 | 0.12 |
| Red/far-red |  |  |  |  |  |  | -0.13 | -0.27 | **0.38** | 0.3 | -0.18 |
| Soil moisture |  |  |  |  |  |  |  | -0.34 | **-0.43** | **0.5** | **0.83** |
| pH |  |  |  |  |  |  |  |  | -0.24 | **-0.4** | -0.15 |
| Soil C/N |  |  |  |  |  |  |  |  |  | -0.13 | **-0.58** |
| N mineral |  |  |  |  |  |  |  |  |  |  | 0.31 |

**Table S4.**
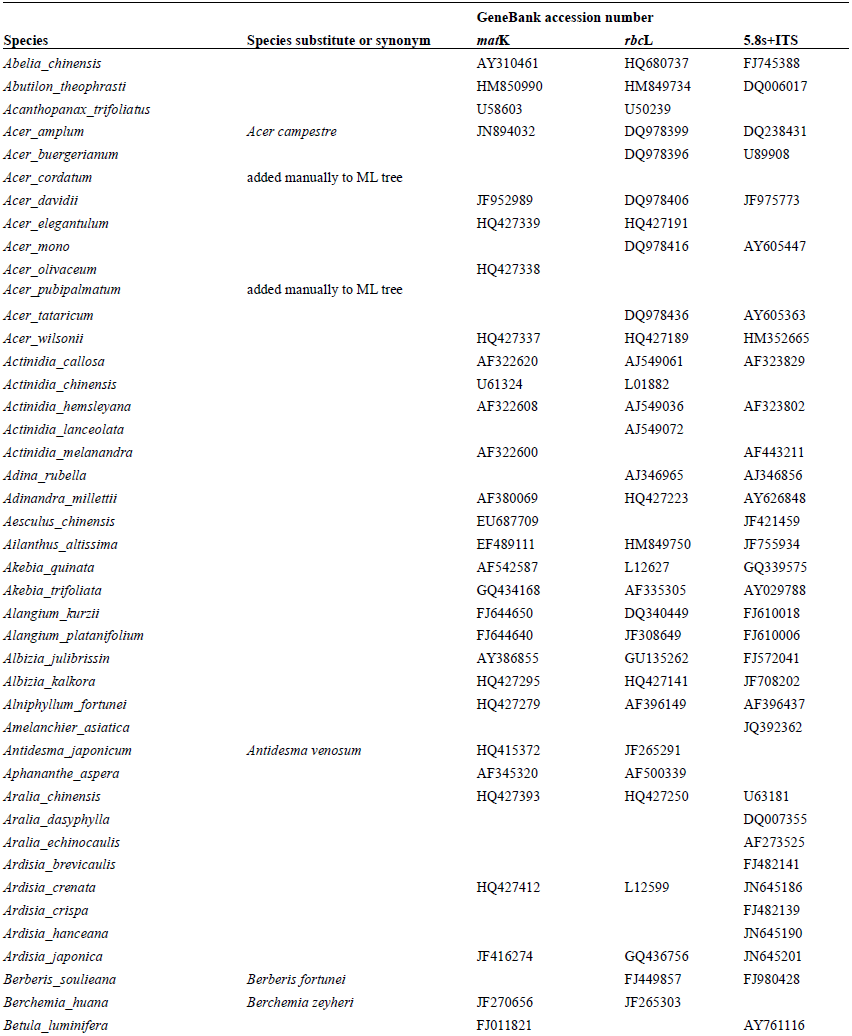
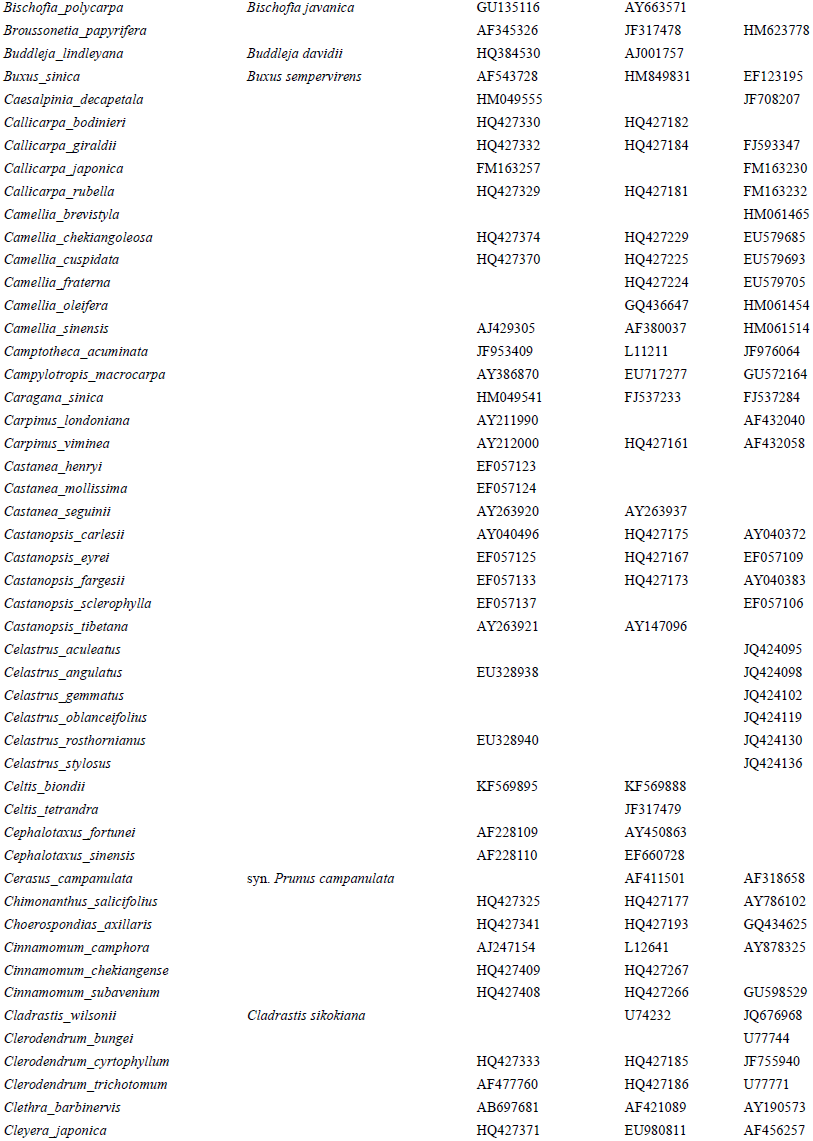
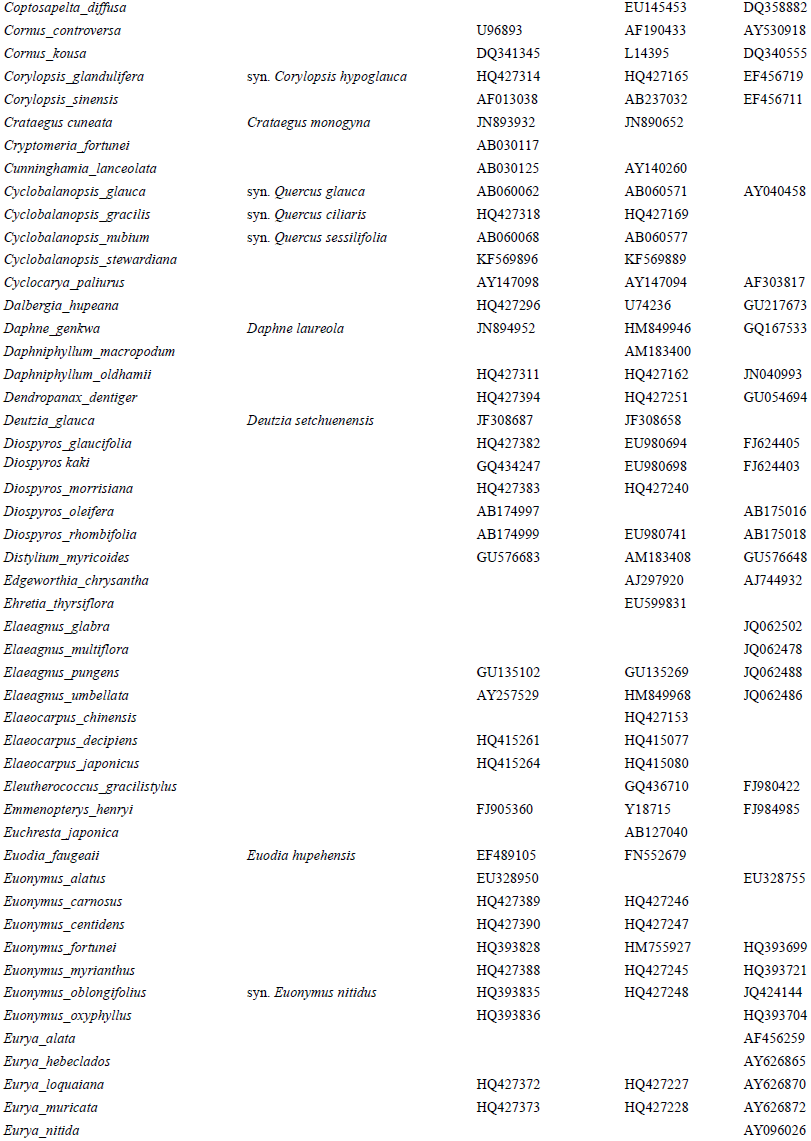
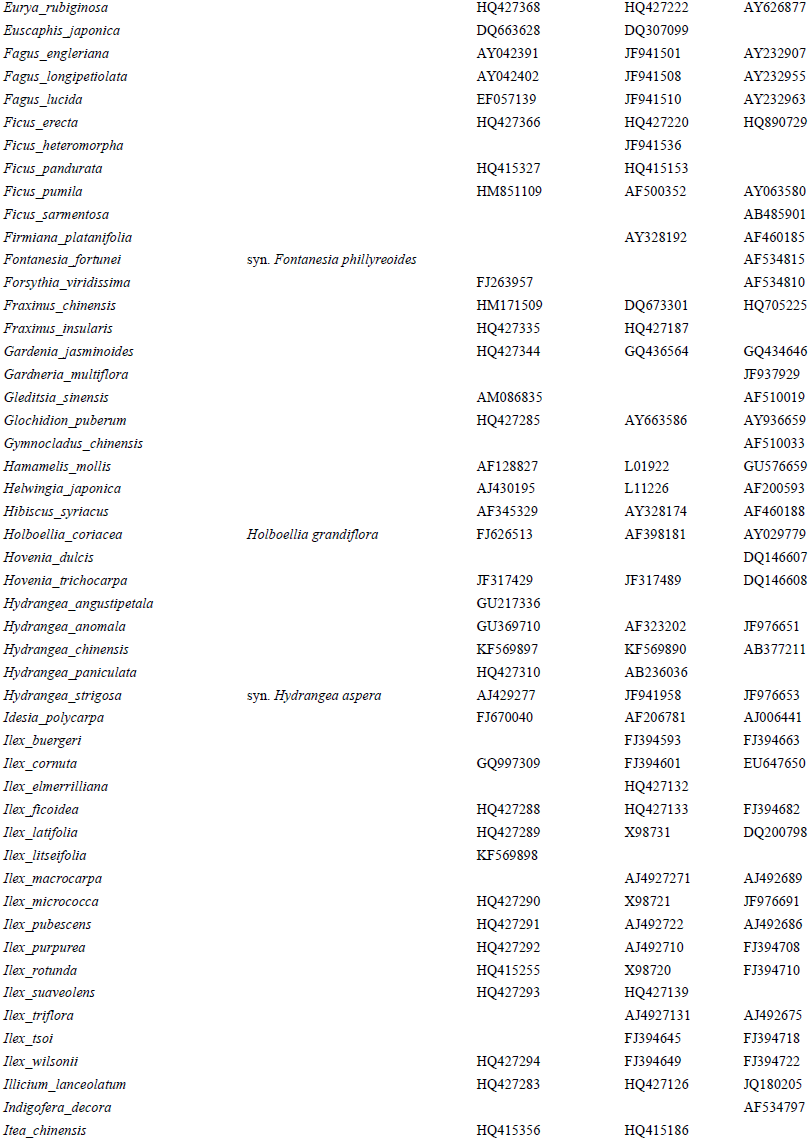
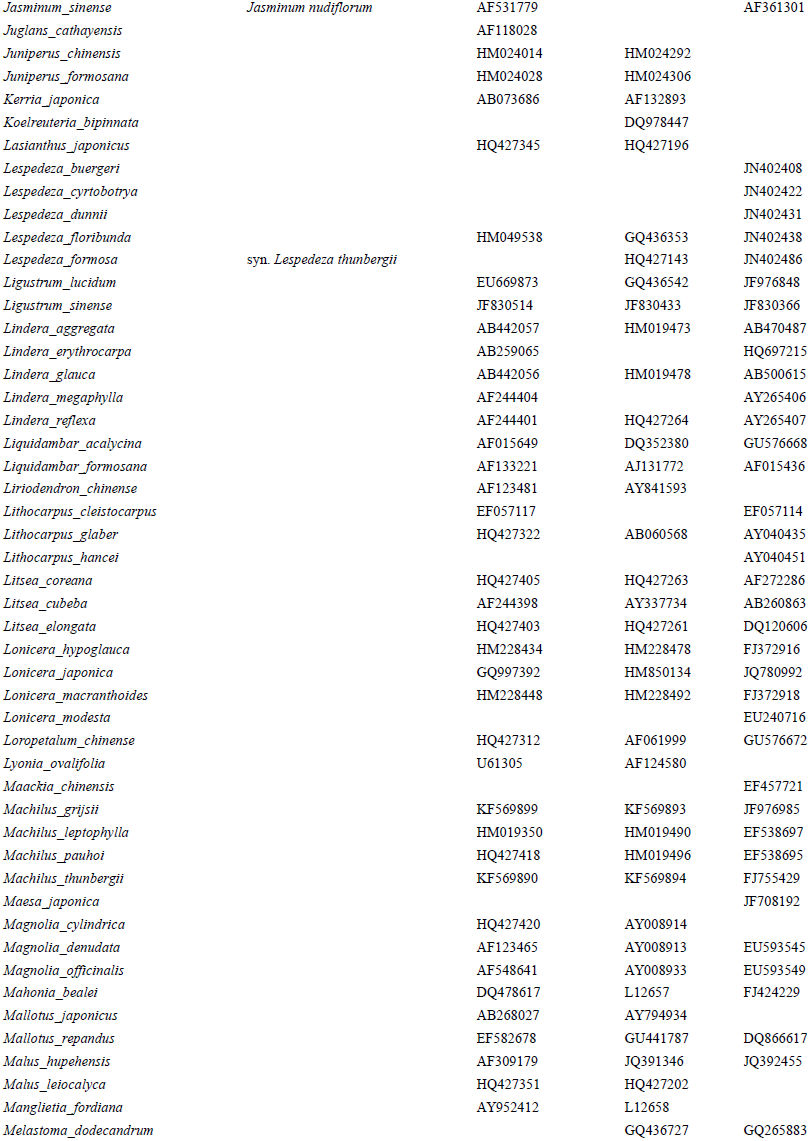
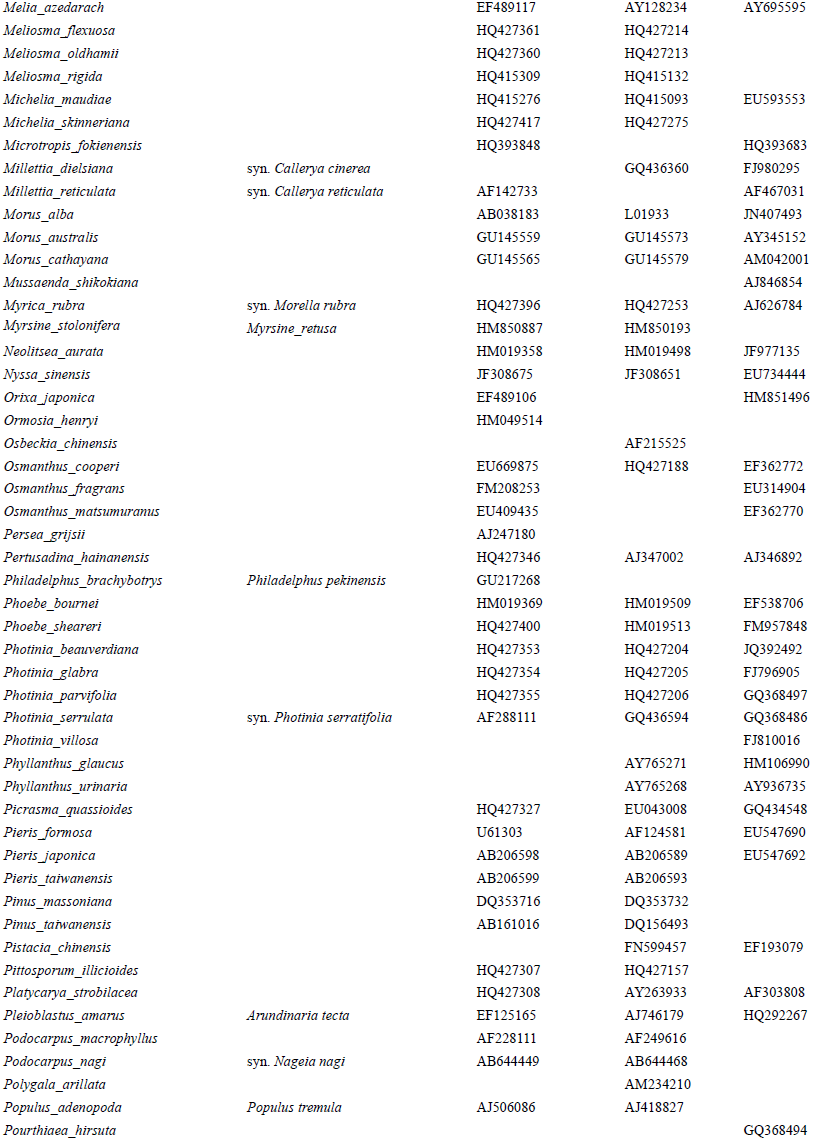
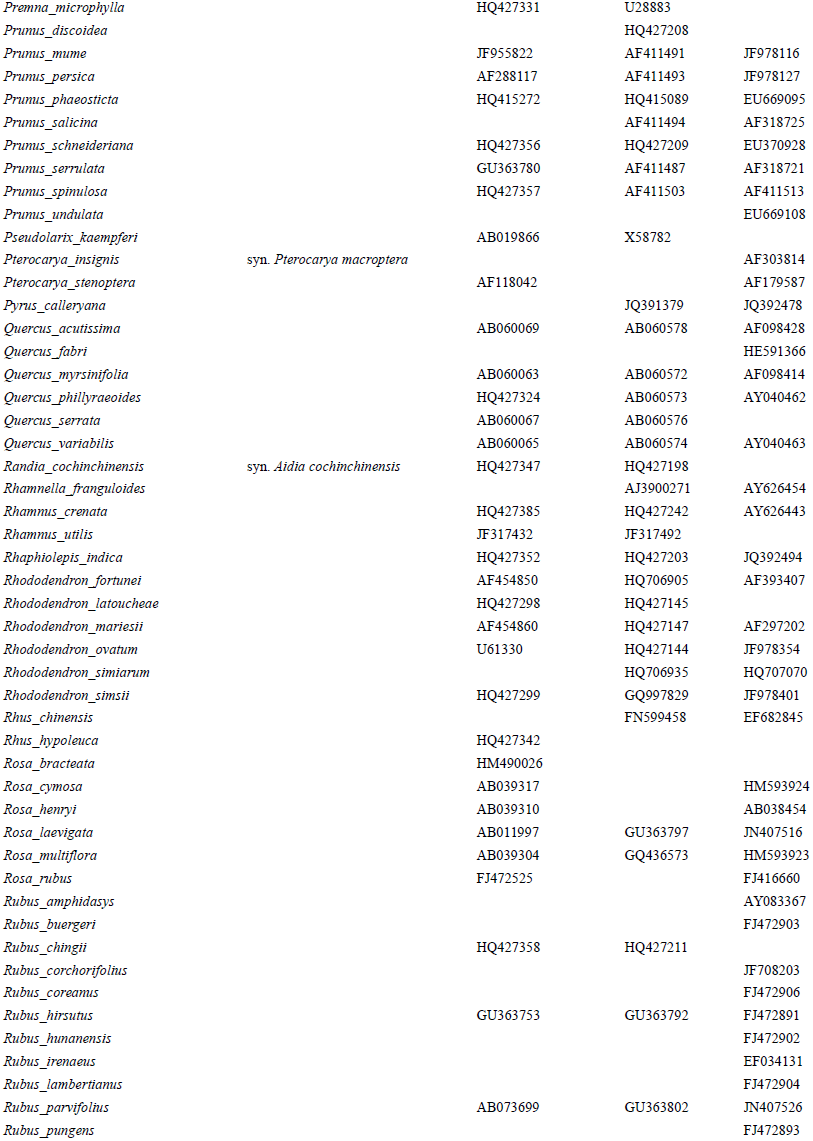
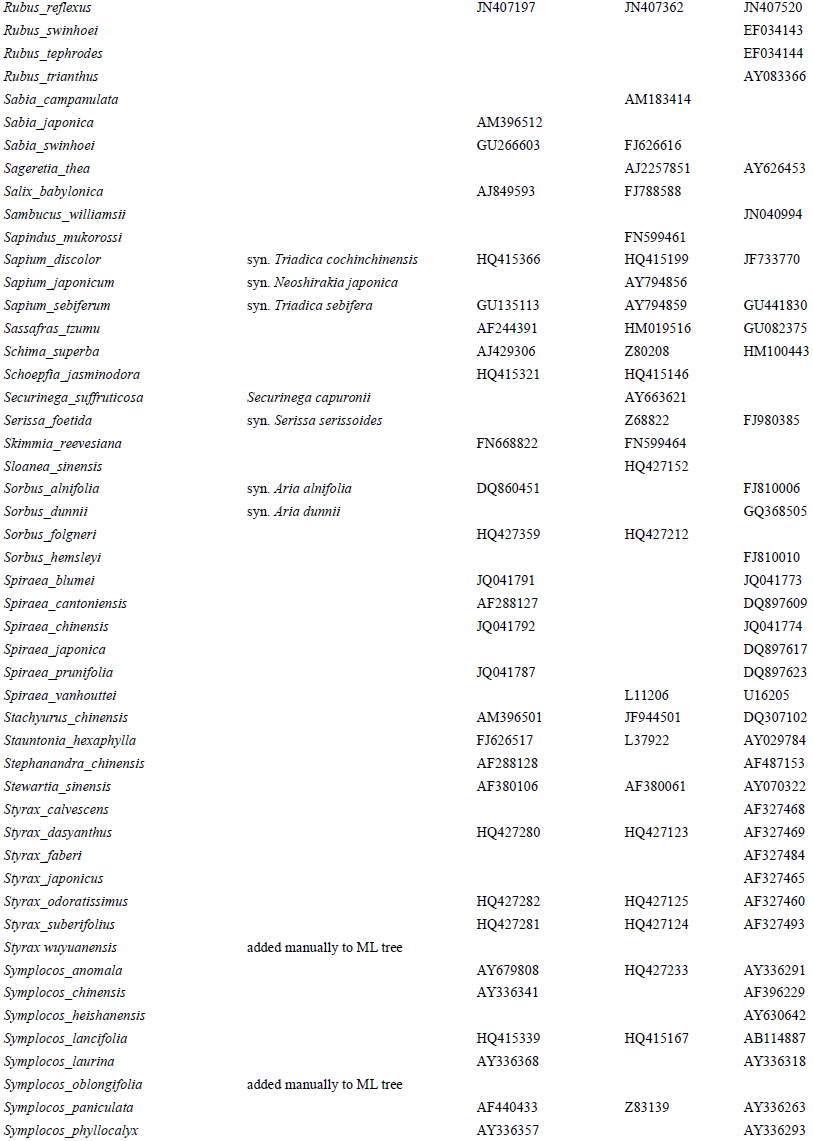
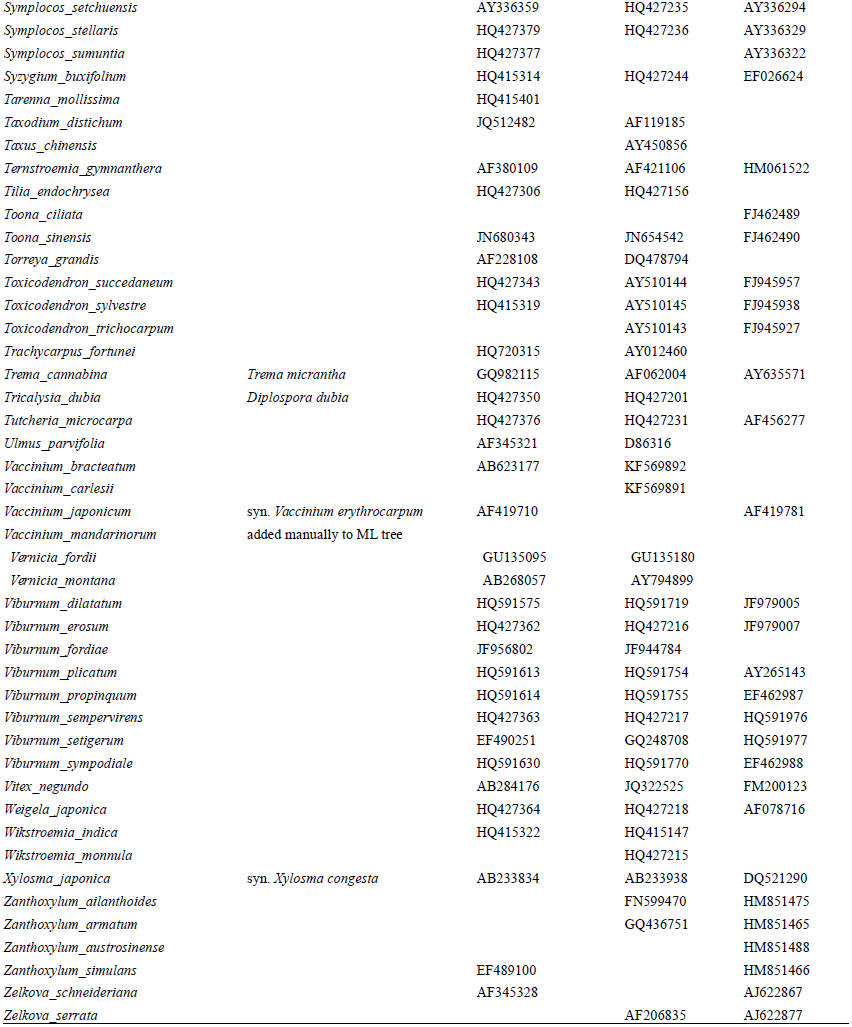
Sequence information for the woody plant species in the Gutianshan National Nature Reserve

**Table S5.** Age constraints for nodes used to create the ultrametric tree.

| Clade | Node defined by MRCA to | Calibration type | Age [ma] | Reference |
| --- | --- | --- | --- | --- |
| Seed plants | *Taxodium distichum* - *Abutilon theophrasti* | max | 385 | (Gerrienne et al. 2004) |
| Gymnosperms | *Pseudolarix kaempferi* - *Taxodium distichum* | min | 318 | (Renner 2009) |
| Cupressaceae | *Cunninghamia lanceolata* - *Taxodium distichum* | min | 90 | (LePage 2003) |
| Pinaceae | *Pseudolarix kaempferi* - *Pinus massoniana* | min | 90 | (Gandolfo et al. 2001)<br>(Hughes and McDougall 1987, |
| Angiosperms | *Pleioblastus amarus* - *Abutilon theophrasti* | max | 130 | Hughes et al. 1991) |
| Laurales | *Chimonanthus salicifolius* - *Litsea cubeba* | min | 108.8 | (Crane et al. 1994) |
| Eudicots | *Holboellia coriacea* - *Abutilon theophrasti* | fixed | 125 | (Hughes and McDougall 1990) |
| Ranunculales | *Holboellia coriacea* - *Mahonia bealei* | min | 91 | (Knobloch and Mai 1986) |
| Berberidaceae | *Berberis soulieana* - *Mahonia bealei* | min | 33.9 | (Manchester 1999)<br>(Magallon-Puebla et al. 1996, |
| Hamamelidaceae | *Liquidambar acalycina* - *Corylopsis sinensis* | min | 83.5 | Magallón et al. 2001) |
| Fabales | *Polygala arillata* - *Albizia kalkora* | min | 60 | (Lavin et al. 2005) |
| Malpighiales | *Vernicia fordii* - *Phyllanthus urinaria* | min | 89.3 | (Crepet and Nixon 1998) |
| Salicaceae | *Idesia polycarpa* - *Populus adenopoda* | min | 48 | (Boucher et al. 2003)<br>(Pactová 1966, Batten 1981, |
| Fagales | *Quercus serrata* - *Juglans cathayensis* | min | 93.5 | Kedves 1989) |
| Juglandaceae | *Cyclocarya paliurus* - *Juglans cathayensis* | min | 55.8 | (Crane et al. 1990) |
| Rosaceae | *Rosa cymosa* - *Prunus pseudocerasus* | min | 37.2 | (Manchester 1999) |
| Ulmaceae | *Ulmus parvifolia* - *Zelkova schneideriana* | min | 33.9 | (Manchester 1999) |
| Rutaceae-Meliaceae | *Melia azedarach* - *Skimmia japonica* | min | 50 | (Corbett and Manchester 2004) |
| Myrtales | *Melastoma dodecandrum* - *Syzygium buxifolium* | min | 60 | (Pigg et al. 1993) |
| Ericales | *Actinidia melanandra* - *Ardisia crenata*<br>*Actinidia melanandra* - *Rhododendron* | min | 89.6 | (Nixon and Crepet 1993) |
| Actinidiaceae (stem node) | *latoucheae* | min | 77.05 | (Schenk and Hufford 2010) |
| Cornaceae | *Alangium kurzii* - *Cornus kousa* | min | 55.8 | (Manchester 1999) |
| Nyssaceae | *Camptotheca acuminata* - *Nyssa sinensis* | min | 33.9 | (Manchester 1999) |
| Hydrangeaceae | *Deutzia glauca* - *Hydrangea strigosa* | min | 37.2 | (Manchester 1999) |
| Cornales | *Camptotheca acuminata* - *Cornus kousa* | min | 89 | (Schenk and Hufford 2010) |
| Oleaceae | *Osmanthus matsumuranus* - *Fraxinus chinensis* | min | 33.9 | (Manchester 1999)<br>(Manchester and Donoghue |
| Dipsacales | *Viburnum sympodiale* - *Lonicera modesta* | min | 33.9 | 1995) |
| Apiales | *Pittosporum illicioides* - *Dendropanax dentiger* | min | 37.2 | (Manchester 1999) |

**Table S6.**
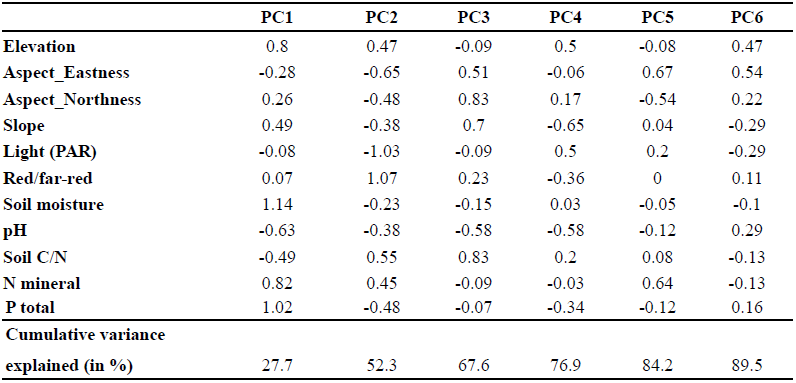
Loadings and percentage of total variation explained of the first six principal components (PCs) of a PCA on the eleven environmental variables. The first two PCs correspond to variation in soil moisture and light, respectively. See Fig. S7 for PCA biplot.

**Table S7.** Phylogenetic signal (Blomberg's *K*, Pagel's *λ* and Abouheif/Moran's *I*) in each of the six traits. Values of Blomberg's *K* and Pagel's *λ* equal to one correspond to a Brownian motion model of trait evolution, while values of *K* or *λ* close to zero indicate no phylogenetic signal. Unlike *K* and *λ,* Abouheif/Moran's *I* is a measure of phylogenetic autocorrelation and is not based on an evolutionary model. *P*-values for the *K-* and *I-* statistics were obtained by randomly shuffling (999 times) the tips on the phyogeny. *P*-values for Pagel's *λ* were obtained based on likelihood-ratio tests.

|  | Leaf area | SLA | Leaf N | Leaf P | Wood density | Height |
| --- | --- | --- | --- | --- | --- | --- |
| Blomberg's $K$ | 0.902 | 0.385 | 0.726 | 0.576 | 0.534 | 0.46 |
| $P$ | < 0.001 | 0.023 | < 0.001 | < 0.001 | < 0.001 | 0.005 |
| Pagel's $\lambda$ | 0.991 | 0.45 | 0.902 | 0.596 | 0.612 | 0.479 |
| $P$ | < 0.001 | < 0.001 | < 0.001 | < 0.001 | < 0.001 | < 0.001 |
| Abouheif/Moran's $I$ | 0.266 | 0.22 | 0.32 | 0.246 | 0.248 | 0.125 |
| $P$ | < 0.001 | < 0.001 | < 0.001 | < 0.001 | < 0.001 | 0.028 |

**Table S8.**
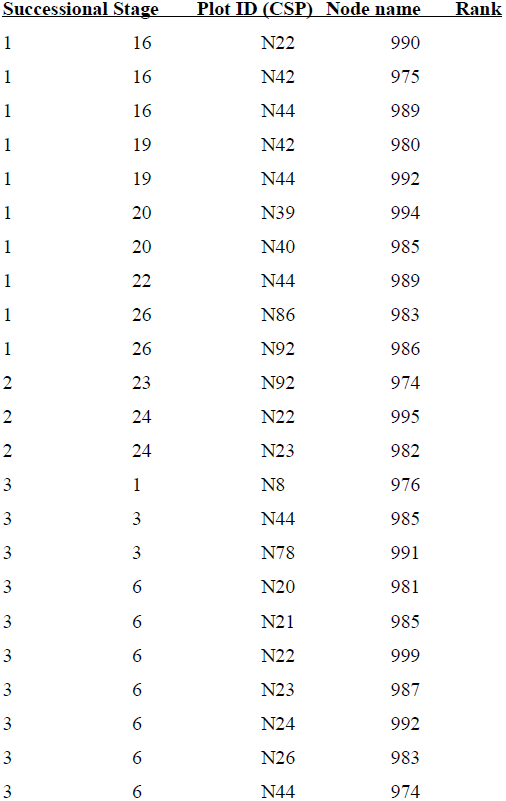
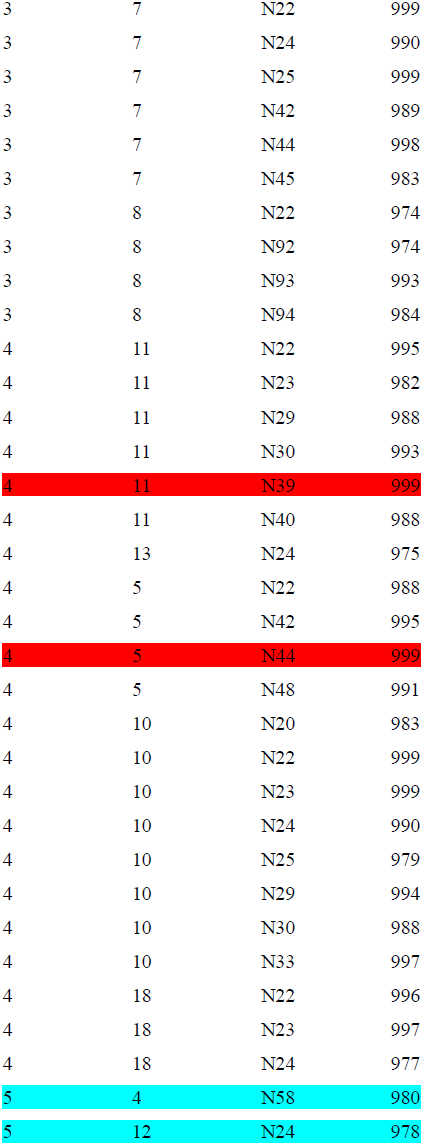
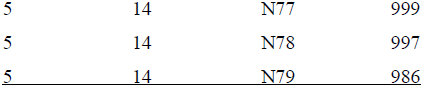
Nodes with significantly more taxa than expected within a particular plot (CSP). Columns correspond to successional stage (Stage 1-5), Plot ID (see also Fig. 1c), node name (see Fig. S6), and ranks in the null distribution across 999 randomization runs, shuffling the tips in the phylogeny. Only nodes that are within the upper 2.5-percentile of the null distribution are listed. Highlighted (for illustration purposes) are the most significant nodes, associated with the plot pairs that had the highest levels of phylogenetic turnover at the two late successional stages (red: stage 4, CSPs 5 & 11; blue: stage 5, CSPs 4 & 12) (see also Fig. 1c and Fig. S6).

**Methods S1** We gathered sequence information, i.e. *mat*K, *rbc*L and the ITS region including the 5.8s gene for all woody species from Gutianshan National Nature Reserve (Lou & Li, 1998) or closely related species available in GenBank (http://www.ncbi.nlm.nih.gov/genbank/, accessed between May and June 2012). For some species of the CSPs, *mat*K and *rbc*L were sequenced using standard barcoding protocols (Fazekas *et al.*, 2012) (Accession numbers: KF569888-KF569899, Table S4). All sequences were aligned separately for the different markers using MAFFT v6 (Katoh *et al.*, 2002).

Sequences for *mat*K and *rbc*L were aligned with the ‘Auto’ option in the online version of the program (http://mafft.cbrc.jp/alignment/server/). The ITS region was aligned with the ‘Q-INS-I’ option considering secondary structure of RNA using the MAFFT application at Bioportal Kumar *et al.*, 2009)). Aligned sequences were concatenated for each species resulting in a total alignment of 3521 nucleotide positions. A phylogenetic tree was inferred using a Maximum Likelihood (ML) method implemented in PhyML (Guindon & Gascuel, 2003). For ML inference, the best fitting model (GTR+I+G) selected by Modeltest (Posada and Crandall 1998) was applied with the following options: tree topology search operation: best of NNI and SPR search, number of substitution rate categories =6, all other parameters were estimated (Gamma Distribution Parameter Alpha, Proportion of Invariable Sites, Transition/Transversion Ratio).

Species occurring in the CSPs but without sequence information available (Table S4) were added manually to the obtained ML tree by the following procedure. *Acer cordatum* was added within *Acer* as a polytomy to the most recent common ancestor (MRCA) of a monophyletic clade formed by other members of *Acer* sect. *Palmata* (i.e. *A*. *elegantulum, A*. *wilsonii, A*. *olivaceum*). Its branch length was defined as the average distance from the MRCA of that clade to the tips. *Styrax wuyuanensis, Symplocos oblongifolia* and *Vaccinium mandarinorum* were added similarily as polytomy emerging from the MRCA for all other members of the respective genus included, with branch lengths equalling the average branch length from that MRCA to the tips of congeners.

Using the ML topology and branch lengths an ultrametric tree was created by non-parametric rate smoothing (nprs) as implemented in r8s (Sanderson, 1997). Absolute node ages were obtained using 27 published fossils or dates as age constrains. A fixed age of 125 million years was applied to the crown node of the Eudicots (Table S5).

**Methods S4** Because non-random phylogenetic structure at the plot scale may simply reflect non-random pattern in overall species frequencies (or abundances) across the phylogeny (Mouquet *et al.*, 2012), we tested for phylogenetic signal in species occurrences as well as abundances at the scale of the whole data set using the APD (abundance phylogenetic deviation) index proposed by Hardy (2008). There was no phylogenetic signal in overall species' occurrence frequencies or abundances in our study (APD = 0.014, *P* = 0.056 and APD = 0.053, *P* = 0.996), so there was no need to implement a null model that restricts permutations to species with similar occurrence frequencies (or abundances).

**Methods 5** We tested, based on 100 simulation runs, whether differences in the number of plots (communities) among stages affect the estimates or phylogenetic turnover (П_ST_ and B_ST_) using the following procedure: in each simulation run we (i) generated 10 communities with 10 species each, and 20 species in total, (ii) calculated П_ST_ (or B_ST_) based on different numbers of plots (3-10 plots) and assessed the Pearson-correlation between П_ST_ (or B_ST_) and the number of plots, and (iii) tested (using a one sample t-test) whether the mean correlation obtained from the 100 simulations significantly differed from zero. Calculations of П_ST_ (or B_ST_) were based on a random Yule (pure-birth) tree for 20 tips [R-package 'phytools' (Revell, 2012)]. We found that the mean correlation between П_ST_ (and B_ST_) and the number plots was close to zero, indicating that there is no intrinsic correlation between the phylogenetic turnover estimates used in our study and the number of plots.

## References

Abouheif E. 1999. A method for testing the assumption of phylogenetic independence in comparative data. Evolutionary Ecology Research 1: 895–909.

Arroyo-Rodríguez V, Melo FPL, Martínez-Ramos M, Bongers F, Chazdon RL, Meave JA, Norden N, Santos BA, Leal IR et al. 2017. Multiple successional pathways in human-modified tropical landscapes: new insights from forest succession, forest fragmentation and landscape ecology research. Biological Reviews 92: 326–340.

Baeten L, Davies TJ, Verheyen K, Van Calster H, Vellend M. 2015. Disentangling dispersal from phylogeny in the colonization capacity of forest understorey plants. Journal of Ecology 103: 175–183.

Baraloto C, Hardy OJ, Paine CET, Dexter KG, Cruaud C, Dunning LT, Gonzalez M-A, Molino J-F, Sabatier D, Savolainen V et al. 2012. Using functional traits and phylogenetic trees to examine the assembly of tropical tree communities. Journal of Ecology 100: 690– 701.

Bartlett MK, Zhang Y, Yang J, Kreidler N, Sun S-W, Lin L, Hu Y-H, Cao K-F, Sack L. 2016. Drought tolerance as a driver of tropical forest assembly: resolving spatial signatures for multiple processes. Ecology 97: 503–514.

Bazzaz FA. 1979. The physiological ecology of plant succession. Annual Review of Ecology and Systematics 10: 351–371.

Bhaskar R, Dawson TE, Balvanera P. 2014. Community assembly and functional diversity along succession post-management. Functional Ecology 28: 1256–1265.

Blomberg SP, Garland Jr T, Ives AR. 2003. Testing for phylogenetic signal in comparative data: behavioral traits are more labile. Evolution 57: 717–745.

Böhnke M, Kreißig N, Kröber W, Fang T, Bruelheide H. 2012. Wood trait-environment relationships in a secondary forest succession in South-East China. Trees 26: 641–651.

Böhnke M, Kröber W, Welk E, Wirth C, Bruelheide H. 2014. Maintenance of constant functional diversity during secondary succession of a subtropical forest in China. Journal of Vegetation Science 25: 897–911.

Bruelheide H, Böhnke M, Both S, Fang T, Assmann T, Baruffol M, Bauhus J, Buscot F, Chen X-Y, Ding B-Y et al. 2011. Community assembly during secondary forest succession in a Chinese subtropical forest. Ecological Monographs 81: 25–41.

Buzzard V, Hulshof CM, Birt T, Violle C, Enquist BJ. 2016. Re-growing a tropical dry forest: functional plant trait composition and community assembly during succession. Functional Ecology 30: 1006–1013.

Cadotte MW. 2014. Including distantly related taxa can bias phylogenetic tests. Proceedings of the National Academy of Sciences 111: E536–E536.

Cadotte MW, Tucker CM. 2017. Should environmental filtering be abandoned? Trends in Ecology & Evolution. 32: 429–437.

Cavender-Bares J, Ackerly DD, Baum DA. Bazzaz FA. 2004. Phylogenetic overdispersion in Floridian oak communities. American Naturalist 163: 823–843.

Cavender-Bares J, Keen A, Miles B. 2006. Phylogenetic structure of Floridian plant communities depends on taxonomic and spatial scale. Ecology 87: 109–122.

Cavender-Bares J, Reich PB. 2012. Shocks to the system: community assembly of the oak savanna in a 40-year fire frequency experiment. Ecology 93: S52–S69.

Chang L-W, Zelený D, Li C-F, Chiu S-T, Hsieh C-F. 2013. Better environmental data may reverse conclusions about niche-and dispersal-based processes in community assembly. Ecology 94: 2145–2151.

Chave J, Coomes D, Jansen S, Lewis SL, Swenson NG, Zanne AE. 2009. Towards a worldwide wood economics spectrum. Ecology Letters 12: 351–366.

Cornell JH, Slatyer RO. 1977. Mechanisms of succession in natural communities and their role in community stability and organization. The American Naturalist 111: 1119–1144.

Ding Y, Zang R, Letcher SG, Liu S, He F. 2012. Disturbance regime changes the trait distribution, phylogenetic structure and community assembly of tropical rain forests. Oikos 121: 1263–1270.

Dornelas M. 2010. Disturbance and change in biodiversity. Philosophical Transactions of the Royal Society B: Biological Sciences 365: 3719–3727.

Du Y, Mi X, Ma K. 2012. Comparison of seed rain and seed limitation between community understory and gaps in a subtropical evergreen forest. Acta Oecologica 44: 11–19.

Eastman JM, Paine CET, Hardy OJ. 2011. spacodiR: structuring of phylogenetic diversity in ecological communities. Bioinformatics 27: 2437–2438.

Eichenberg D, Purschke O, Ristok C, Wessjohann L, Bruelheide H. 2015. Trade-offs between physical and chemical carbon-based leaf defence: of intraspecific variation and trait evolution. Journal of Ecology 103: 1667–1679.

Fine PVA, Kembel SW. 2011. Phylogenetic community structure and phylogenetic turnover across space and edaphic gradients in western Amazonian tree communities. Ecography 34: 552–565.

Fukami T, Martijn Bezemer T, Mortimer SR, van der Putten WH. 2005. Species divergence and trait convergence in experimental plant community assembly. Ecology Letters 8: 1283–1290.

Garnier E, Cortez J, Billes R.s G, Navas M-L, Roumet C, Debussche M, Laurent G, Blanchard A, Aubry D, Bellmann A et al. 2004. Plant functional markers capture ecosystem properties during secondary succession. Ecology 85: 2630–2637.

Graham CH, Macháč A, Storch D. 2016. Phylogenetic scale in ecology and evolution. bioRxiv 063560.

Hardy OJ. 2008. Testing the spatial phylogenetic structure of local communities: statistical performances of different null models and test statistics on a locally neutral community. Journal of Ecology 96: 914–926.

Hardy OJ. 2010. SPACoDi 0.10: a program for spatial and phylogenetic analysis of community diversity. [WWW document] URL http://ebe.ulb.ac.be/ebe/Software.html. [accessed 7 February 2013].

Hardy OJ, Couteron P, Munoz F, Ramesh BR, Pélissier R. 2012. Phylogenetic turnover in tropical tree communities: impact of environmental filtering, biogeography and mesoclimatic niche conservatism. Global Ecology & Biogeography 21: 1007–1016.

Hardy OJ, Senterre B. 2007. Characterizing the phylogenetic structure of communities by an additive partitioning of phylogenetic diversity. Journal of Ecology 95: 493–506.

Helmus MR, Bland TJ, Williams CK, Ives AR. 2007. Phylogenetic measures of biodiversity. The American Naturalist 169: E68–E83.

Hu Z, Yu M. 2008. Study on successions sequence of evergreen broad-leaved forest in Gutian Mountain of Zhejiang, Eastern China: species diversity. Frontiers of Biology in China 3: 45–49.

Jombart T, Balloux F, Dray S. 2010. Adephylo: new tools for investigating the phylogenetic signal in biological traits. Bioinformatics 26: 1907–1909.

Keddy PA. 1992. Assembly and response rules – 2 goals for predictive community ecology. Journal of Vegetation Science 3: 157–164.

Kröber W, Böhnke M, Welk E, Wirth C, Bruelheide H et al. 2012. Leaf trait-environment relationships in a subtropical broadleaved forest in South-east China. PLoS One 7: e35742.

Kunstler G, Lavergne S, Courbaud B, Thuiller W, Vieilledent G, Zimmermann NE, Kattge J, Coomes DA. 2012. Competitive interactions between forest trees are driven by species' trait hierarchy, not phylogenetic or functional similarity: implications for forest community assembly. Ecology Letters 15: 831–840.

Legendre P, Mi XC, Ren HB, Ma KP, Yu MJ, Sun IF, He FL. 2009. Partitioning beta diversity in a subtropical broad-leaved forest of China. Ecology 90: 663–674.

Letcher SG. 2010. Phylogenetic structure of angiosperm communities during tropical forest succession. Proceedings of the Royal Society B: Biological Sciences 277: 97–104.

Letcher SG, Chazdon RL, Andrade AC, Bongers F, van Breugel M, Finegan B, Laurance SG, Mesquita RC, Martínez-Ramos M, Williamson GB. 2012. Phylogenetic community structure during succession: evidence from three Neotropical forest sites. Perspectives in Plant Ecology, Evolution and Systematics 14: 79–87.

Letcher SG, Lasky JR, Chazdon RL, Norden N, Wright SJ, Meave JA, Pérez-García EA, Muñoz R, Romero-Pérez E, Andrade A et al. 2015. Environmental gradients and the evolution of successional habitat specialization: a test case with 14 Neotropical forest sites. Journal of Ecology 103: 1276–1290

Letten AD, Keith DA, Tozer MG. 2014. Phylogenetic and functional dissimilarity does not increase during temporal heathland succession. Proceedings of the Royal Society of London B: Biological Sciences 281: 20142102.

Levin SA, Muller-Landau HC. 2000. The evolution of dispersal and seed size in plant communities. Evolutionary Ecology Research 2: 409–435.

Li SP, Cadotte MW, Meiners SJ, Hua ZS, Jiang L, Shu WS. 2015. Species colonisation, not competitive exclusion, drives community overdispersion over long-term succession. Ecology Letters 18: 964–973.

Losos JB. 2008. Phylogenetic niche conservatism, phylogenetic signal and the relationship between phylogenetic relatedness and ecological similarity among species. Ecology Letters 11: 995–1003.

Lou LH, Li GY. 1988. Biota of Gutianshan Nature Reserve: list of seed plants. [In Chinese] Manuscript.

Lozupone CA, Hamady M, Kelley ST, Knight R. 2007. Quantitative and qualitative beta diversity measures lead to different insights into factors that structure microbial communities. Applied and Environmental Microbiology 73: 1576–1585.

Magurran A. 2004. Measuring biological diversity. Oxford, UK, Blackwell Publishing.

Mayfield MM, Levine JM. 2010. Opposing effects of competitive exclusion on the phylogenetic structure of communities. Ecology Letters 13: 1085–1093.

Moles AT, Warton DI, Warman L, Swenson NG, Laffan SW, Zanne AE, Pitman A, Hemmings FA, Leishman MR. 2009. Global patterns in plant height. Journal of Ecology 97: 923–932.

Mouquet N, Devictor V, Meynard CN, Munoz F, Bersier L-F, Chave J, Couteron P, Dalecky A, Fontaine C, Gravel D et al. 2012. Ecophylogenetics: advances and perspectives. Biological Reviews 87: 769–785.

Norden N, Letcher SG, Boukili V, Swenson NG, Chazdon R. 2012. Demographic drivers of successional changes in phylogenetic structure across life-history stages in plant communities. Ecology 93: S70–S82.

Oksanen J, Blanchet FG, Friendly M, Kindt R, Legendre P, McGlinn D, Minchin PR, O’Hara RB, Simpson GL, Solymos P et al. 2017. vegan: community ecology package. R package ver. 2.4-3 [WWW document] URL http://r-forge.r-project.org/projects/vegan/. [accessed 7 April 2017].

Pagel M. 1999. Inferring the historical patterns of biological evolution. Nature 401: 877–884.

Parmentier I, Hardy OJ. 2009. The impact of ecological differentiation and dispersal limitation on species turnover and phylogenetic structure of inselberg's plant communities. Ecography 32: 613–622.

Parmentier I, Réjou-Méchain M, Chave J, Vleminckx J, Thomas DW, Kenfack D, Chuyong GB, Hardy OJ. 2014. Prevalence of phylogenetic clustering at multiple scales in an African rain forest tree community. Journal of Ecology 102: 1008–1016.

Pavoine S, Bonsall MB. 2011. Measuring biodiversity to explain community assembly: a unified approach. Biological Reviews 86: 792–812.

Preston FW. 1960. Time and space and the variation of species. Ecology 41: 611–627.

Purschke O, Schmid BC, Sykes MT, Poschlod P, Michalski SG, Durka W, Kühn I, Winter M, Prentice HC. 2013. Contrasting changes in taxonomic, phylogenetic and functional diversity during a long-term succession: insights into assembly processes. Journal of Ecology 101: 857–866.

Purschke O, Sykes MT, Poschlod P, Michalski SG, Römermann C, Durka W, Kühn I, Prentice HC. 2014. Interactive effects of landscape history and current management on dispersal trait diversity in grassland plant communities. Journal of Ecology 102: 437–446.

R Development Core Team. 2017. R: a language and environment for statistical computing. R foundation for statistical computing.

Revell LJ. 2012. phytools: an R package for phylogenetic comparative biology (and other things). Methods in Ecology and Evolution 3: 217–223.

Roeder M, McLeish M, Beckschäfer P, de Blécourt M, Paudel E, Harrison RD, Slik F. 2015. Phylogenetic clustering increases with succession for lianas in a Chinese tropical montane rain forest. Ecography 38: 832–841.

Shipley B, Vile D, Garnier E. 2006. From plant traits to plant communities: a statistical mechanistic approach to biodiversity. Science 314: 812–814.

Swenson NG, Enquist BJ, Thompson J, Zimmerman JK. 2007. The influence of spatial and size scale on phylogenetic relatedness in tropical forest communities. Ecology 88: 1770–1780.

Swenson NG, Stegen JC, Davies SJ, Erickson DL, Forero-Montana J, Hurlbert AH, Kress WJ, Thompson J, Uriarte M, Wright SJ et al. 2012. Temporal turnover in the composition of tropical tree communities: functional determinism and phylogenetic stochasticity. Ecology 93: 490–499.

Swenson NG. 2013. The assembly of tropical tree communities–the advances and shortcomings of phylogenetic and functional trait analyses. Ecography 36: 264–276.

Uriarte M, Swenson NG, Chazdon RL, Comita LS, Kress WJ, Erickson D, Forero-Montaña J, Zimmerman JK, Thompson J. 2010. Trait similarity, shared ancestry and the structure of neighbourhood interactions in a subtropical wet forest: implications for community assembly. Ecology Letters 13: 1503–1514.

Vellend M, Cornwell W, Magnuson-Ford K, Mooers A. 2011. Measuring phylogenetic biodiversity. In: Magurran A, McGill B, eds. Biological diversity: frontiers in measurement and assessment. Oxford, UK: Oxford University Press, 197–207.

Wang X-H, Kent M, Fang X-F. 2007. Evergreen broad-leaved forest in Eastern China: its ecology and conservation and the importance of resprouting in forest restoration. Forest Ecology and Management 245: 76–87.

Webb C, Ackerly D, Kembel S. 2009. Phylocom. Software for the analysis of phylogenetic community structure and character evolution. ver. 4.2. [WWW document] URL http://phylodiversity.net/phylocom/. [accessed: 16 December 2012].

Westoby M, Falster DS, Moles AT, Vesk PA, Wright IJ. 2002. Plant ecological strategies: some leading dimensions of variation between species. Annual Review of Ecology and Systematics 33: 125–159.

Wheeler B, Torchiano M. 2016. lmPerm: permutation tests for linear models. R package ver. 2.1.0. [WWW document] URL http://cran.r-project.org/web/packages/lmPerm/. [accessed: 18 August 2016].

White EP, Ernest SKM, Adler PB, Hurlbert AH, Lyons SK. 2010. Integrating spatial and temporal approaches to understanding species richness. Philosophical Transactions of the Royal Society B: Biological Sciences 365: 3621–3631.

Whitfeld TJS, Kress WJ, Erickson DL, Weiblen GD. 2012. Change in community phylogenetic structure during tropical forest succession: evidence from New Guinea. Ecography 35: 821–830.

Wright IJ, Reich PB, Westoby M, Ackerly DD, Baruch Z, Bongers F, Cavender-Bares J, Chapin T, Cornelissen JH, Flexas J. 2004. The worldwide leaf economics spectrum. Nature 428: 821–827.

Wu Z, others. 1980. Vegetation of China. [In Chinese.] Science Press Beijing China.

## References

Batten DJ. 1981. Stratigraphic, palaeogeographic and evolutionary significance of late cretaceous and early tertiary normapolles pollen. Review of Palaeobotany and Palynology 35: 125–137.

Boucher LD, Manchester SR, Judd WS. 2003. An extinct genus of Salicaceae based on twigs with attached flowers, fruits, and foliage from the Eocene Green River Formation of Utah and Colorado, USA. American Journal of Botany 90: 1389–1399.

Corbett SL, Manchester SR. 2004. Phytogeography and fossil history of Ailanthus (Simaroubaceae). International Journal of Plant Sciences 165: 671–690.

Crane PR, Friis EM, Pedersen KR. 1994. Palaeobotanical evidence on the early radiation of magnoliid angiosperms. Plant Systematics and Evolution - Supplementa 8: 51–72.

Crane PR, Manchester SR, Dilcher DL. 1990. A preliminary survey of fossil leaves and well-preserved reproductive structures from the Sentinel Butte Formation (Paleocene) near Almont, North Dakota. Fieldiana. Geology 20: 1–63.

Crepet WL, Nixon KC. 1998. Fossil Clusiaceae from the Late Cretaceous (Turonian) of New

Jersey and implications regarding the history of bee pollination. American Journal of Botany 85: 1122–1133.

Fazekas AJ, Kuzmina ML, Newmaster SG, Hollingsworth PM. 2012. DNA Barcoding Methods for Land Plants In: DNA Barcodes: Methods and protocols (eds. Kress WJ, Erickson DL), pp. 223–252. Humana Press, New York.

Gandolfo MA, Nixon KC, Crepet WL. 2001. Turonian Pinaceae of the Raritan Formation, New Jersey. Plant Systematics and Evolution 226: 187–203.

Gerrienne P, Meyer-Berthaud B, Fairon-Demaret M, Streel M and Steemans P. 2004. Runcaria, a Middle Devonian Seed Plant Precursor. Science 306: 856–858.

Guindon S, Gascuel O. 2003. A Simple, Fast, and Accurate Algorithm to Estimate Large Phylogenies by Maximum Likelihood. Systematic Biology 52: 696–704.

Hughes NF, McDougall AB. 1987. Records of angiospermid pollen entry into the english early cretaceous succession. Review of Palaeobotany and Palynology 50: 255–272.

Hughes NF, McDougall AB. 1990. Barremian-Aptian angiospermid pollen records from southern England. Review of Palaeobotany and Palynology 65: 145–151.

Hughes NF, McDougall AB, Chapman JL. 1991. Exceptional new record of Cretaceous Hauterivian angiospermid pollen from Southern England. Journal of Micropalaeontology 10: 75–82.

Katoh K, Misawa K, Kuma K, Miyata, T. 2002. MAFFT: a novel method for rapid multiple sequence alignment based on fast Fourier transform. Nucleic Acids Research 30: 3059–3066.

Kedves M. 1989. Evolution of the Normapolles complex. In: Evolution, Systematics, and Fossil History of the Hamamelidae, 1-7. Systematics Association Special Volume, vol. 40B. (eds. Crane P. R, Blackmore S.). Clarendon Press, Oxford.

Knobloch E, Mai DH. 1986. Monographie der Früchte und Samen in der Kreide von Mitteleuropa, Praha.

Kumar S, Skjaeveland A, Orr R, Enger P, Ruden T, Mevik B-H, Burki F, Botnen A, Shalchian-Tabrizi K. 2009. AIR: A batch-oriented web program package for construction of supermatrices ready for phylogenomic analyses. BMC Bioinformatics 10: 357.

Lavin M, Herendeen PS, Wojciechowski MF. 2005. Evolutionary rates analysis of Leguminosae implicates a rapid diversification of lineages during the Tertiary. Systematic Biology 54: 575–594.

LePage BA. 2003. The evolution, biogeography and palaeoecology of the Pinaceae based on fossil and extant representatives. Acta Horticulturae 615: 29–52.

Lou LH, Li GY. 1998. List of seed plants in Gutianshan.

Magallon-Puebla S, Herendeen PS, Endress PK. 1996. Allonia decandra: Floral remains of the tribe Hamamelideae (Hamamelidaceae) from Campanian strata of southeastern USA. Plant Systematics and Evolution 202: 177–198.

Magallón S, Herendeen PS, Crane P. 2001. Androdecidua endressii gen. et sp. nov, from the Late Cretaceous of Georgia (United States): Further Floral Diversity in Hamamelidoideae (Hamamelidaceae). International Journal of Plant Sciences 162: 963–983.

Manchester SR. 1999. Biogeographical relationships of North American tertiary floras. Annals of the Missouri Botanical Garden 86: 472–522.

Manchester SR, Donoghue MJ. 1995. Winged fruits of Linnaeeae (Caprifoliaceae) in the Tertiary of Western North America: Diplodipelta gen. nov. International Journal of Plant Sciences 156: 709–722.

Nixon KC, Crepet WL. 1993. Late Cretaceous fossil flowers of ericalean affinity. American Journal of Botany 80: 616–623.

Pacltová B. 1966. Pollen grains of angiosperms in the Cenomanian Peruc Formation in Bohemia. Palaeobotanist 15: 52–54.

Pigg KB, Stockey RA, Maxwell SL. 1993. Paleomyrtinaea, a new genus of permineralized myrtaceous fruits and seeds from the Eocene of British Columbia and Paleocene of North Dakota. Canadian Journal of Botany 71: 1–9.

Posada D, Crandall KA. 1998. MODELTEST: testing the model of DNA substitution. Bioinformatics 14: 817–818.

Renner S. 2009. Gymnosperms. In: The Timetree of Life (eds. Hedges SB, Kumar S), pp. 157–160. Oxford University Press, Oxford.

Sanderson MJ. 1997. A nonparametric approach to estimating divergence times in the absence of rate constancy. Molecular Biology and Evolution 14: 1218–1231.

Schenk JJ, Hufford L. 2010. Effects of substitution models on divergence time estimates:

Simulations and an empirical study of model uncertainty using Cornales. Systematic Botany 35: 578–592.

